# Spatial and temporal requirement of Mlp60A isoforms during muscle development and function in *Drosophila melanogaster*

**DOI:** 10.1101/2021.12.18.473287

**Authors:** Rohan Wishard, Mohan Jayaram, S. R. Ramesh, Upendra Nongthomba

## Abstract

Many myofibrillar proteins undergo isoform switching in a spatio-temporal manner during muscle development. The biological significance of the variants of several of these myofibrillar proteins remains elusive. One such myofibrillar protein, the <u>M</u>uscle <u>L</u>IM <u>P</u>rotein (MLP), is a vital component of the Z-discs. In this paper, we show that one of the Drosophila MLP encoding genes, *Mlp60A*, gives rise to two isoforms: a short (279 bp, 10 kDa) and a long (1461 bp, 54 kDa) one. The short isoform is expressed throughout development, but the long isoform is adult-specific, being the dominant of the two isoforms in the indirect flight muscles (IFMs). A concomitant, muscle-specific knockdown of both isoforms leads to late pupal lethality, with the surviving flies being majorly flight defective. *Mlp60A* null flies show developmental lethality, and muscle defects in the individuals surviving till the third instar larval stage. This lethality could be rescued partially by muscle-specific overexpression of the short isoform. Almost 90% of the long isoform-specific *P-element* insertion mutant flies show a compromised flight ability and have reduced sarcomere length. Hence, our data shows that the two Mlp60A isoforms are functionally specialized, to ensuring normal embryonic muscle development and adult flight muscle function.

## Introduction

The postnatal development of vertebrate cardiac and skeletal muscles is marked by switching of several sarcomeric contractile proteins from their foetal to respective adult isoforms [1–7]. However, the cellular mechanisms by which these isoforms are regulated during striated muscle development still have not been studied in detail. The first step towards understanding this phenomenon is to elucidate the functional significance of this developmental isoform switching [8], and the redundancy/non-redundancy among the different isoforms of several sarcomeric contractile proteins [1, 9–11]. The mixed population of different pure and hybrid fibre types in mammalian skeletal muscles makes it difficult to study the functions of the specific isoforms of sarcomeric proteins [4, 12]. On the other hand, the *Drosophila* dorsal longitudinal muscles (DLMs), which are a type of indirect flight muscles (IFMs), offer a unique model system to study muscle development, due to their structural similarity to the vertebrate skeletal muscles [13, 14], a functional similarity with the vertebrate cardiac muscles [15, 16], and a relatively simpler composition of only ‘fibrillar’ type fibres [16, 17]. Moreover, the later stages of DLM development mimic the postnatal development of vertebrate striated muscles with regards to isoform switching of sarcomere proteins. Several sarcomeric proteins such as Myosin Heavy Chain (MHC), Actin, Troponin subunits, Tropomyosin, Myosin Light Chain (MLC), Kettin and Zormin, etc., are known to undergo isoform switching from their embryonic/larval to their respective adult isoforms, that are either expressed specifically in the DLMs, or both the DLMs and the jump muscle-TDT (Tergal Depressor of Trochanter) [18–26]. Consequently, DLM- or DLM-TDT-specific null mutants of different sarcomeric proteins have been isolated, which facilitate the study of these stage-specific isoforms, and the importance of maintaining the correct isoforms in specific muscles during development [11, 22, 25, 27–30]. In the present study, we have shown that the <u>M</u>uscle <u>L</u>IM <u>P</u>rotein at <u>60A</u> (Mlp60A) undergoes isoform switching during muscle development in *Drosophila*. The vertebrate ortholog of Mlp60A is CSRP3. Cell culture studies have demonstrated that this protein can promote myogenic differentiation by associating with several muscle-specific transcription factors, such as MyoD, MRF4, and myogenin [31–32]. Moreover, through studies performed in a knockout mouse model, this protein has been shown to be necessary for the development of cardiomyocyte cytoarchitecture. In fact, *CSRP3^−/−^* mice show both dilated cardiomyopathy (DCM)- and hypertrophic cardiomyopathy (HCM)-like phenotypes [33, 34]. *CSRP3* mutations have also been identified in human cardiomyopathy patients [35, 36, 37, 38]. However, the precise role of MLP in muscle differentiation, and its requirement for skeletal muscle development have not been addressed. MLP is expressed in both developing and adult skeletal musculature in both mice and zebrafish, but its complete deficiency in either of these model organisms produces very mild skeletal muscle phenotypes [33, 39–40]. Hence, it is not understood if this protein is dispensable for skeletal muscle development, or whether some alternate isoform or paralog compensate for its deficiency. Interestingly, an alternate isoform of MLP, called the “MLP-b” isoform, was reported by Vafiadaki *et al*., which was found to be upregulated in tissue samples from skeletal muscle disease patients, and appeared to be a negative regulator of myogenesis [41]. However, the distinct spatio-temporal requirements of the two MLP isoforms in skeletal muscle development, and the precise role of the novel MLP-b isoform were not addressed. Our results show that there is an exclusive functional specialization of the *Mlp60A* isoforms in *Drosophila*, with one isoform being constitutive, and essential for embryonic muscle development, and the other isoform being adult-specific, and necessary for the development of myofibrils with normal sarcomere length and function in the IFMs.

## Results

### The *Mlp60A* locus in *Drosophila melanogaster* codes for two isoforms, resulting from alternative splicing, with distinct spatio-temporal expression profiles, which localize to the Z-discs of the sarcomeres

The *Mlp60A* locus is predicted to produce a putative, second, alternatively spliced isoform (Fig. 1A, https://flybase.org/reports/FBgn0259209.html#gene_model_products), in addition to the isoform reported earlier [42]. We were able to experimentally validate the presence of both these isoforms in cDNA prepared from the whole body (Fig. 1B). The respective full-length amplicons of both the isoforms were sequenced and analysed (Fig. S1A). The sequence of the long isoform CDS was submitted to GenBank (NCBI Accession No: MN990115). The sequence alignment revealed that the two isoforms share a common transcription start site and the first two exons of the locus. An alternative splicing event, as shown in Fig. S1B, leads to expression of the long isoform. To check whether the long isoform is indeed translated, we generated polyclonal antibodies against the short isoform CDS (common to both isoforms) (Fig. S2). Using the resulting serum, we were able to detect a bigger, 54 kDa isoform, corresponding to the long transcript (1461 bp) and a smaller, 10 kDa isoform, corresponding to the short transcript (279 bp) (Fig. 1B-C).

**Figure 1:**
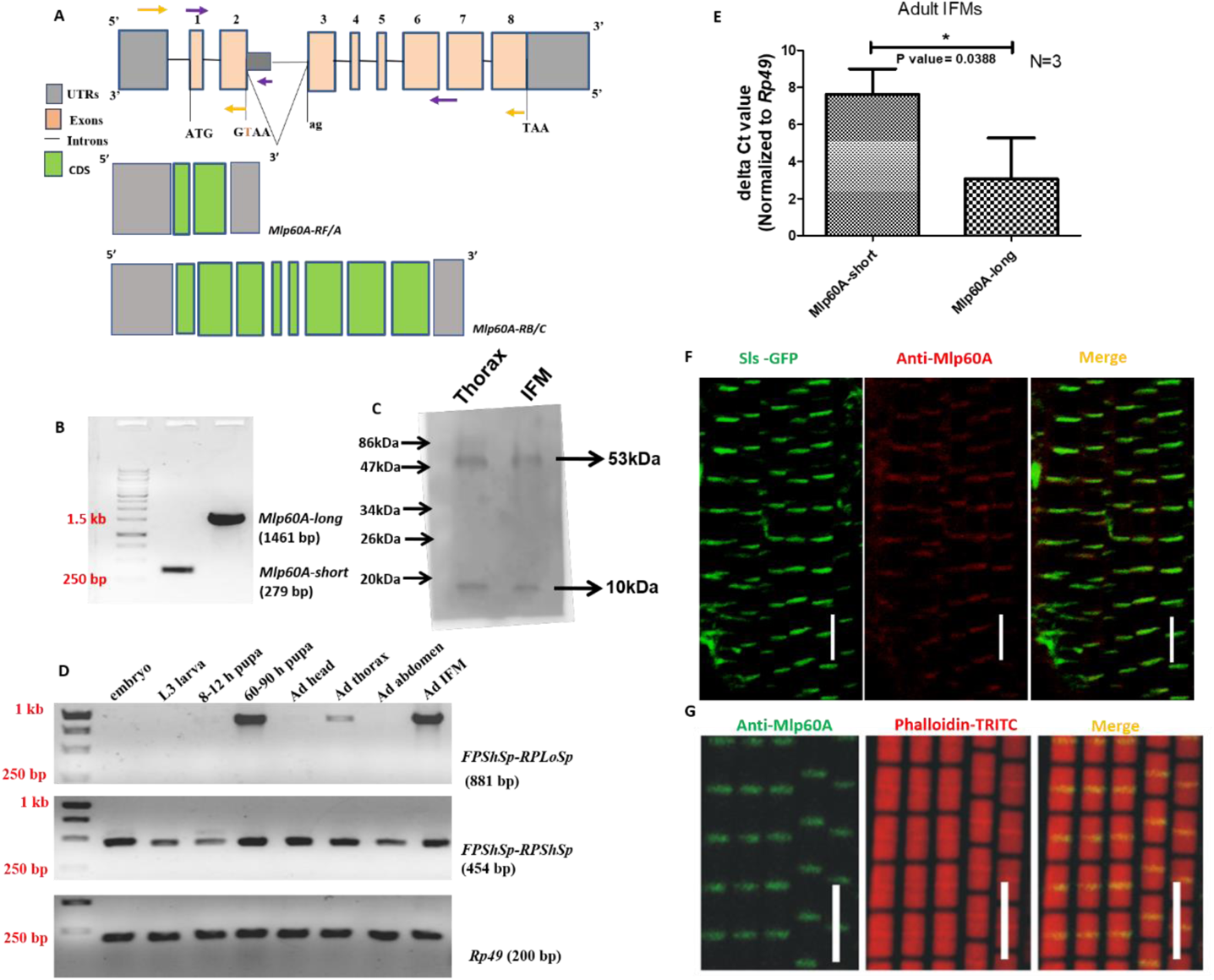
Expression and developmental switching of alternatively spliced isoforms of Mlp60A. (A) Gene locus with the experimentally verified short isoform (*Mlp60A-RF/A*) and the bioinformatically predicted long isoform (*Mlp60A-RB/C*). (B) 1% Agarose Gel showing the full length *Mlp60A-short* and *Mlp60A-long* CDS, amplified from newly eclosed adult whole-body cDNA. These were amplified using primer combinations shown in yellow. A common forward primer: *FPShIsoCl*, was used with two different reverse primers: *RPShIsoCl* for the short isoform and *RPLoIsoCl* for the long isoform. (C) Western blot showing the two isoforms, detected with Mlp60A polyclonal antibody. (D) Expression profiling of *Mlp60A-short* and *Mlp60A*-*long* isoforms by qualitative RT-PCR. These were amplified using primer combinations shown in purple. A common forward primer (*FPShSp*) was used with a short isoform-specific (*RPShSp*) or long isoform-specific (*RPLoSp*) reverse primer, to perform this experiment. (E) Quantitative analysis of *Mlp60A-short* (using *Mlp60A_exon2_FP*/*RPShSp* primer pair) and *Mlp60A-long* (using *FPLoSp*/*RPLoSp* primer pair) isoforms by qRT-PCR. (F) Immunostaining of adult IFMs with anti-Mlp60A polyclonal antibody, in Sls-GFP background, strongly co-localize to Z-discs. (G) Co-immunostaining of adult IFMs with anti-Mlp60A polyclonal antibodies and Phalloidin-TRITC. Scale bar reads 5 micro-meters (µm).

The expression pattern of the two isoforms was analysed across different developmental stages and in various tissues of the adult. As shown in Fig. 1D, the short isoform is expressed constitutively across all the developmental stages, and in different body segments of the adult. However, the long isoform expression commences only during the mid-pupal stages, following which it becomes restricted to the thorax, being very prominent in the IFMs. Since the IFMs express both the isoforms, their expression was analysed quantitatively in the adult IFMs. As shown in Fig. 1E, in the adult IFMs, the expression of the long isoform was found significantly higher than that of the short isoform. Immunostaining of IFMs with Mlp60A polyclonal antibodies, either in a *sallimus (sls)-GFP* background (Fig. 1F) or with Phalloidin-TRITC co-immunostaining (Fig. 1G), revealed that the Mlp60A antibodies localize specifically to the Z-discs of the sarcomeres with no signal detected from elsewhere in the cytoplasm. Hence this result shows that both the isoforms localize to the sarcomere Z-discs in the IFMs.

### Concomitant muscle-specific knockdown of both the Mlp60A isoforms leads to pupal lethality and severe flight defect in the surviving progeny

To dissect the developmental and functional requirement of *Mlp60A* in *Drosophila,* we performed a conditional knockdown of both isoforms, beginning from late-embryo/early L1 stage, using the *Gal4/UAS* system [43]. In order to restrict the knockdown to specifically in the muscles, the *Dmef2-Gal4* line was used to drive two different RNAi lines for *Mlp60A* (both containing shRNA targeting the constitutive 2^nd^ exon). Around 86% (Fig. 2A) and 94% (Fig. 2B) knockdown was achieved with RNAi lines 1 and 2, respectively. As shown in Fig. 2C, in both cases, a substantial number of pupae failed to eclose. However, the severity of this phenotype varied depending on the RNAi line used. Where, in case of RNAi line 2-mediated knockdown, as many as 72% of the pupae showed lethality, in case of knockdown using RNAi line 1, the pupal lethality observed was 51%. As shown in Fig. 2D, although the knockdown pupae failed to eclose, they did survive up till the late pupal stages. Furthermore, the knockdown individuals that did survive till the adult stage were tested for their flight ability. As shown in Fig. 2E, 87% of the RNAi line 1-mediated surviving knockdown flies were flight-defective, wherein 47% of flies had defective flight when knocked down with RNAi line 2.

**Figure 2:**
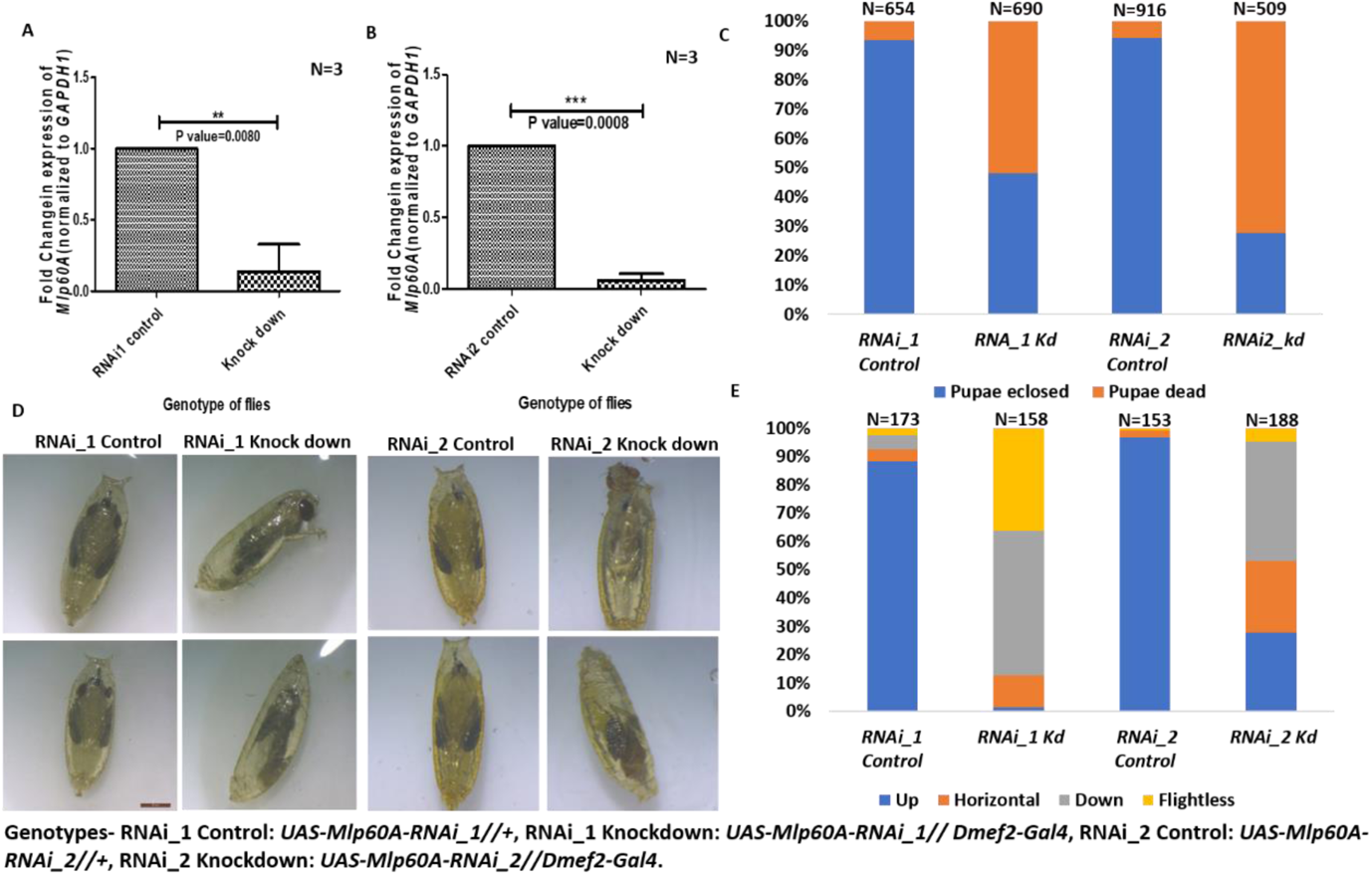
Phenotypic effects of *Dmef2-Gal4* mediated knockdown of both *Mlp60A* isoforms. (A-B) Validation of knockdown with RNAi lines 1 (A) and 2 (B). (C) Assessment of pupal lethality in *Mlp60A* knockdown flies. Y-axis shows the percentage of pupae, X-axis shows the genotype of pupae. (D) Images of dead *Mlp60A* knockdown pupae. The scale bar reads 200 nm. (E) Flight ability of *Mlp60A* knockdown flies. Y-axis shows the percentage of flies, X-axis shows the genotype of flies.

### *Mlp60A* knockdown flies with defective flight show myofibrillar defects

In order to study the muscle phenotypes resulting from knockdown of both Mlp60A isoforms, the flight-defective *Dmef2-Gal4::UAS-Mlp60A-RNAi_1* (knockdown) flies were processed for visualizing the IFM patterning and fascicular structure using polarized optics. As shown in Fig. S3, the IFMs knocked down flies did not differ significantly from the control IFMs in either their pattern (Fig. S3A) or fascicular dimensions (Fig. S3B-C). Next, other flies of the same genotype were processed for observing the IFM myofibrillar structure using confocal microscopy. As shown in Fig. 3A, while the IFMs of the knocked down flies mostly showed myofibrils which were comparable in appearance to those of the control, they did contain a few frayed myofibrils (represented by white arrows). Also, the knockdown IFMs had a reduced resting sarcomere length as compared to the control ones (see a magnified view of the longitudinal sections in Fig. 3A, quantification shown in Fig. 3D). The cross-sections of *Mlp60A*-knockdown IFMs revealed several actin-rich aggregates, which, although majorly concentrated along the IFM fascicular membrane (represented by white arrows in Fig. 3B), were also seen on the membranes of the internal myofibres (represented by black arrows in Fig. 3B). Most importantly, this phenotype was completely penetrant in the *Mlp60A*-knockdown flies, which was not observed in any of their control counterparts. This phenotype suggests that the Mlp60A could be involved in the regulation of thin filament assembly and actin dynamics. There was no significant difference in the fascicular cross-sectional area between the control and knockdown flies (Fig. 3C).

**Figure 3:**
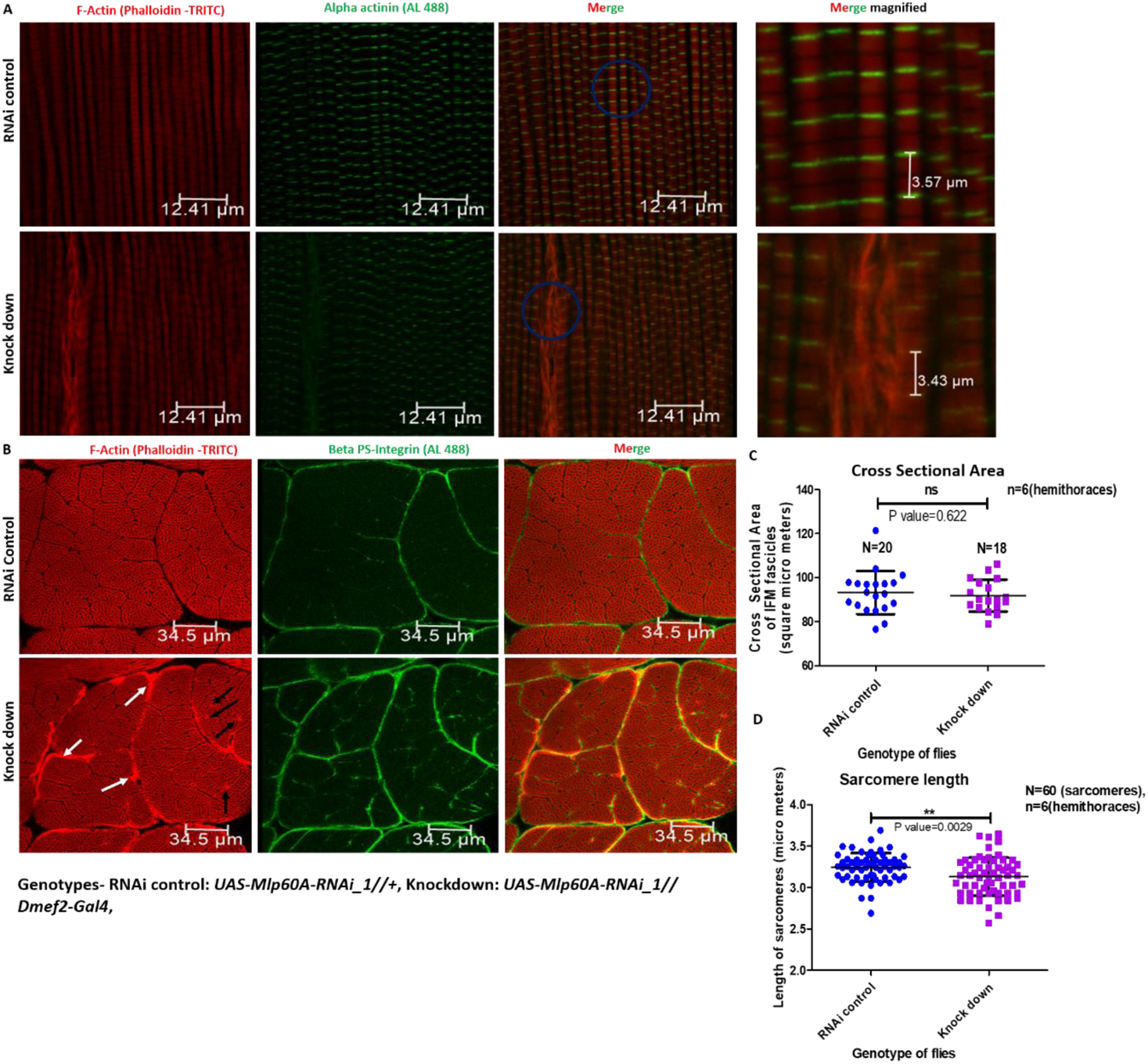
IFM myofibrillar defects in *Mlp60A* knockdown flies. (A) Longitudinal sections of IFMs from *Mlp60A* knocked down and the corresponding control flies, stained for F-actin filaments (red channel) and for *α*-actinin (green channel). The frayed myofibrils seen in *Mlp60A* knocked down flies is marked with white arrows. (B) Transverse (Cross) sections of IFMs from *Mlp60A* knocked down and the corresponding control flies, stained for F-actin filaments (red channel) and *β*-PS Integrin (green channel). Actin rich aggregates in the knocked down IFMs, deposited on the fascicular membrane and the membrane of internal myofibres, are marked with white and black arrows, respectively. (C) Quantification of cross-sectional area of IFM fascicles. (D) Quantification of sarcomere length of IFM myofibrils. Genotypes-RNAi_1 Control: *UAS-Mlp60A-RNAi_1(III)*, RNAi_1 Knockdown: *Dmef2-Gal4(III)::UAS-Mlp60A-RNAi_1(III)*.

### Isolation and characterization of *Mlp60A* isoform-specific mutant alleles by *P-element* hop out mutagenesis

To study the respective functions of the two isoforms, we performed a hop out mutagenesis screen to isolate isoform-specific mutant alleles of *Mlp60A*. Flies having the *P-element* insertion (confirmed as shown in Fig. S4) were crossed with flies coding for the delta 2-3 transposase, to obtain the hop out progeny in the next generation (described in Materials and Methods). We screened for only recessive alleles which, in homozygous condition, caused developmental lethality and/or weak or defective flight in the surviving flies or both (Table 1), consistent with the phenotypes observed in the knockdown experiments.

In order to characterize the mutant alleles, we began by selecting a suitable control allele, which would be one resulting from the precise excision of the inserted *P-element* and the 8 bp target site duplication [44], which a *P-element* insertion is known to cause, thus restoring the sequence at the insertion site to the wild type sequence. For this purpose, several candidate alleles which did not give rise to any developmental lethality, when homozygous, were further tested (Fig. S5A). As shown in Fig. S5A, *UR 2.2.10* homozygous flies flew almost as good as the wild type reference. PCR amplification and sequencing of the region flanking the insertion site, from these flies (Fig. S5B), confirmed that this particular allele was indeed a precise excision allele. Hence, the allele *UR 2.2.10* was renamed as the ‘PEC’ (precise excision control) allele and will be referred to by this name, further in the manuscript.

Having selected a suitable control allele, we proceeded to characterize the respective genomic lesions in each of the homozygous lethal alleles, by performing PCR using overlapping primer pairs covering the insertion site (Fig. S6A). As shown in Fig. S6B-D), the lethal allele *UR 2.9.17*, was found to lack genomic region spanning the first two exons of the *Mlp60A* region, the 5’-UTR, the intergenic region between *Mlp60A* and its upstream gene, *CG3209,* as well as the last two exons of *CG3209*. The allele *UR 2.10.7* was found to harbour a remnant of the *P-element* insertion at the original insertion site (Fig. S6B, lane *UR 2.10.7*). Individuals homozygous for each of these alleles were further characterized for the *Mlp60A* transcripts produced by them, and the phenotype of the corresponding homozygous mutants have been characterized further in this manuscript. The allele *UR 2.4.1* was found to be lacking a portion of the intergenic region between exons 2 and 3 of the *Mlp60A* region (Fig. S6B, lane *UR 2.4.1*). This allele was later found to complement both, the *Mlp60A^null^* allele (characterized in detail in this report) and a genomic deficiency covering the *Mlp60A* region, with regards to survival and flight ability (Fig. S13). Hence this allele was not characterized further.

### Loss of *Mlp60A* leads to severe developmental lethality and larval body wall muscle defects

*UR 2.9.17* homozygous L3 larvae were analysed for the presence of any *Mlp60A* transcript, by RT-PCR. As shown in Fig. 4A, the *Mlp60A* short transcript (the only isoform detected at this stage in the wild type flies, Fig. 1D) could not be detected, even when the reaction was carried out up to saturation. Henceforth, the *UR 2.9.17* allele will be referred to as the *Mlp60A^null^* allele. To determine the effective lethal phase of the homozygous *Mlp60A^null^* individuals, we performed a lethality test, beginning from the L1 stage. As shown in Fig. 4B, most of the null individuals perished during the larval stage itself. As a readout for muscle defects, *Mlp60A^null^* L3 larvae were assayed for their crawling ability. As shown in Fig. 4C-D, the homozygous *Mlp60A^null^* L3 larvae crawled significantly lesser distances compared to the homozygous *PEC* larvae, when tested for the same duration of time. Some of these larvae were then dissected to observe the body wall muscles. As shown in Fig. 4E-F, the *Mlp60A^null^* homozygous larvae showed defective body wall muscles. They showed significant thinning of the body wall muscles, as compared to the control larvae (Fig. 4F). While most of the null larvae had degenerated or malformed muscles, few showed a more severe ‘missing muscles’ phenotype. Fig. 4E shows two representative images of the larval body wall muscles from each of the genotypes tested (defective/degenerated or absent muscles have been marked with white arrows).

**Figure 4:**
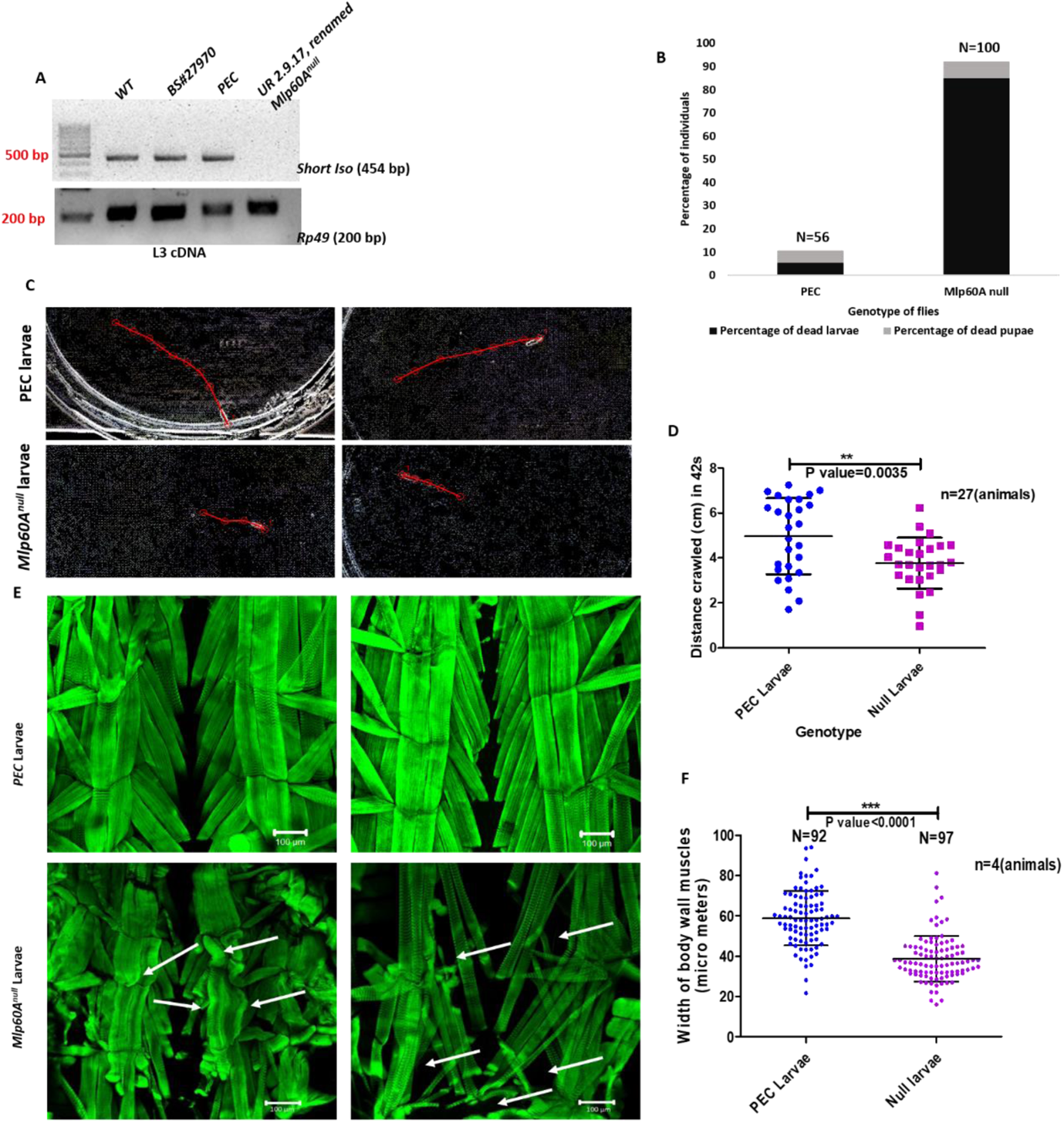
Phenotype of *Mlp60A^null^* homozygous larvae. (A) Analysis of *Mlp60A-short* expression in *UR 2.9.17* homozygous, BS27970# and *PEC* homozygous L3 larvae, along with *w^1118^* larvae as control. Following this, the allele *UR 2.9.17* was renamed as *Mlp60A^null^* allele. (B) Assessment of developmental lethality of *Mlp60A^null^* homozygous larvae, beginning from the L1 developmental stage. (C) Two representative traces, each for *Mlp60A^null^* homozygous and *PEC* homozygous L3 larvae, on 1 % agar plate. (D) Quantification of crawling ability of *Mlp60A^null^* homozygous L3 larvae. (E) Body wall muscle defects visible in *Mlp60A^null^* L3 larvae. The malformed muscles and regions of missing muscles have been marked with white arrows. (F) Quantification of the width of body wall muscles in *Mlp60A^null^* homozygous L3 larvae.

To confirm that it is the abrogation of *Mlp60A* expression which is solely responsible for the phenotypes observed in the *Mlp60A^null^* homozygous larvae, we performed a complementation test between the *Mlp60A^null^* allele and a genomic deficiency covering the *Mlp60A* region: *Df(2R)BSC356* (verified in Fig. S7B). The majority of the *Mlp60A^null^//Df(2R)BSC356* progeny also failed to complete development (Fig. S7A). Not only were the L3 heterozygotes found to have severely compromised crawling ability (Fig. S7C-D), but featured body wall muscle defects reminiscent of those observed in the *Mlp60A^null^* homozygotes (compare Fig. S7E and F to Fig. 4E and F, respectively).

### Muscle-specific transgenic expression of Mlp60A short isoform partially rescues the developmental lethality associated with the *Mlp60A^null^* mutation

In order to test whether the developmental lethality of Mlp60A null mutants is specifically due to the absence of the Mlp60A-short isoform, and to check whether the short isoform alone can compensate for the absence of the long isoform in adults, we sought to perform a rescue experiment by expressing only the Mlp60A short isoform in the *Mlp60A^null^* background. For this purpose, we generated a transgenic fly line in which *UAS* regulatory sequences drive the expression of the full-length *Mlp60A-short* isoform (see Materials and Methods, and Fig. S8-9). *UH3-Gal4* driver line, which has mild ubiquitous expression, including muscles, during development [45], was used to achieve an appreciable overexpression (Fig S10A). The overexpression, by itself alone, did not elicit any phenotype (Fig. S10B). As shown in Fig. 5A, ectopic expression of the short isoform driven by *UH3-Gal4,* showed partial rescue of the developmental lethality observed in the *Mlp60A^null^* mutants. The conditional expression level of the *Mlp60A-short* transcript was lesser than its endogenous expression levels in the wild type, as shown in Fig. 5B-C. The rescued flies were also tested for their flight ability. As shown in Fig. 5D, around 40% of the rescued flies were flight defective, whereas another 20% showed reduced flight ability, being horizontally flighted. This flight profile was drastically different than that of the corresponding positive control flies, 80% of which were Up flighted. These results show that the short isoform alone cannot completely compensate for the loss of the long isoform, in the adults.

**Figure 5:**
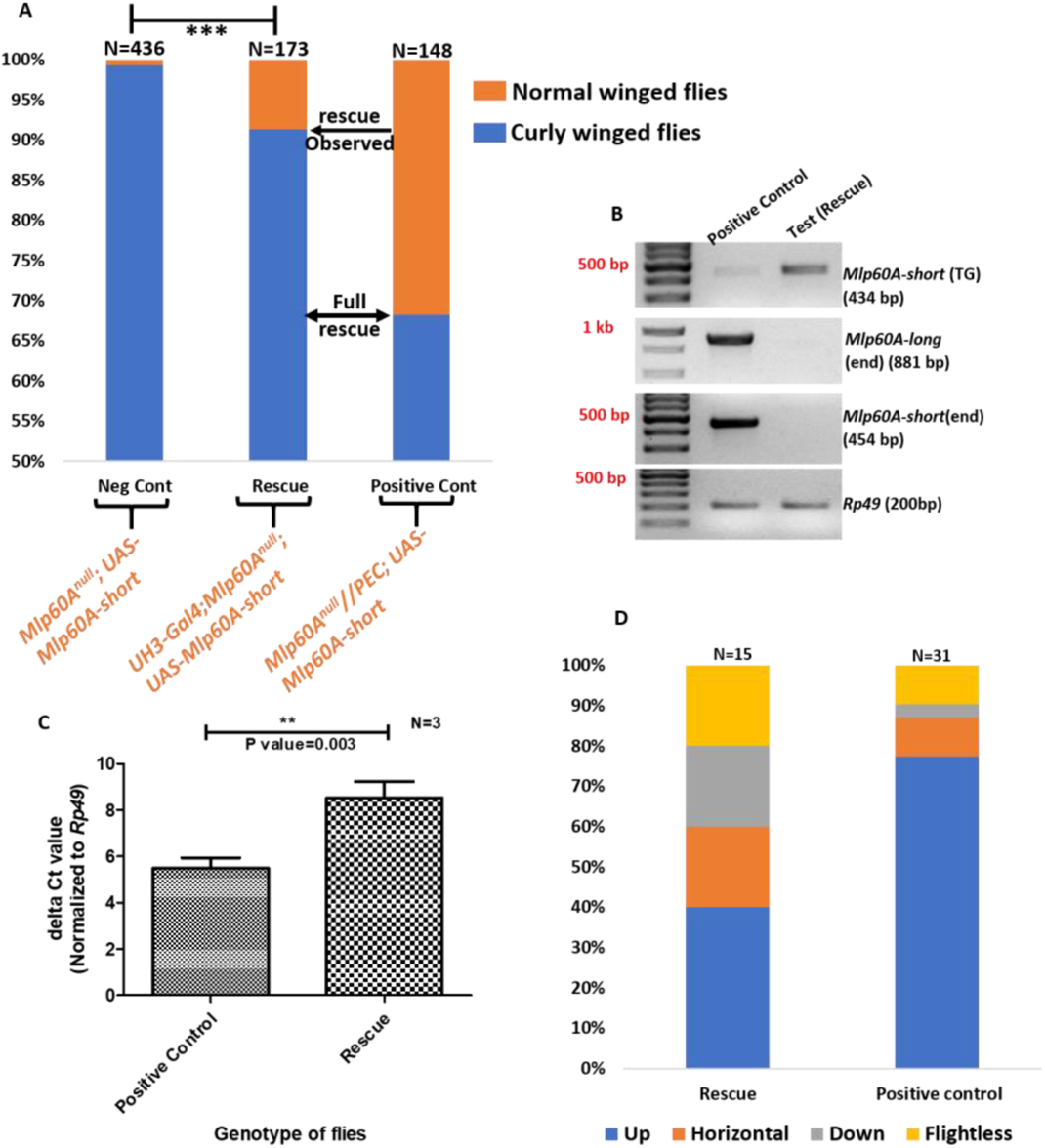
Rescue of Mlp60A null individuals by transgenic overexpression of *Mlp60A-short* isoform. (A) Shows the percentage of normal and curly winged flies (Y-axis) obtained in the test (rescue), negative and positive control sets (X-axis). For difference in the percentage of normal winged flies in the negative control and test sets, P value<0.0001, Chi Squared Value (CSV)=64.646, df=1. A complete rescue would have yielded a percentage of normal winged flies, comparable to that in the positive control (shown by the double headed arrow). The single headed arrow shows the percentage of rescue observed. Genotypes of normal winged flies in each set have been specified in orange font. For detailed methodology, see ‘Materials and Methods’ (B) Shows the qualitative analysis of endogenous *Mlp60A-short*, *Mlp60A-long* and transgenic *Mlp60A-short* isoforms from positive control and rescued fly-thoraces. (C) Shows the quantitative estimation of the levels of endogenous *Mlp60A-short* and transgenic *Mlp60A-short* from the positive control and rescued flies, respectively. (D) Shows the flight ability of the rescued and positive control flies. Y-axis shows the percentage of flies, X-axis shows the genotype of flies. Genotypes-Negative control: *Mlp60A^null^; UAS-Mlp60A-short//+*; Positive control: *PEC//Mlp60A^null^; UAS-Mlp60A-short//+*; Rescue (test): *UH3-Gal4//+; Mlp60A^nulll^; UAS-Mlp60A-short//+*.

### Loss of Mlp60A leads to down-regulation of several thin filament proteins

We hypothesised that loss of Mlp60A could lead to down-regulation of other thin and/or thick filament proteins. We based this hypothesis on three lines of reasoning. First, it is already known from the literature that loss of any specific muscle contractile protein results in a co-ordinated transcriptional down-regulation of other contractile proteins [24, 30, 46–47]. Second, the mammalian ortholog of *Mlp60A*, *CSRP3*, is necessary for muscle differentiation [31–32]. Third, our own results revealed that *Mlp60A^null^* larvae possess thinner muscle fibres (Fig. 4E and F). Hence, to test our hypothesis, we analysed the expression of six major muscle contractile proteins, in *Mlp60A^null^* mutant larvae. Interestingly, expression of the majority of thin filament proteins was found to be significantly down-regulated in *Mlp60A^null^* homozygotes as compared to the control (Fig. 6). Myosin heavy chain (MyHC), the only thick filament protein analysed, did not vary significantly between the test and the control larvae. This could be one of the reasons for thin and defective myofibrils seen in null embryos and larva (discussed later).

**Figure 6:**
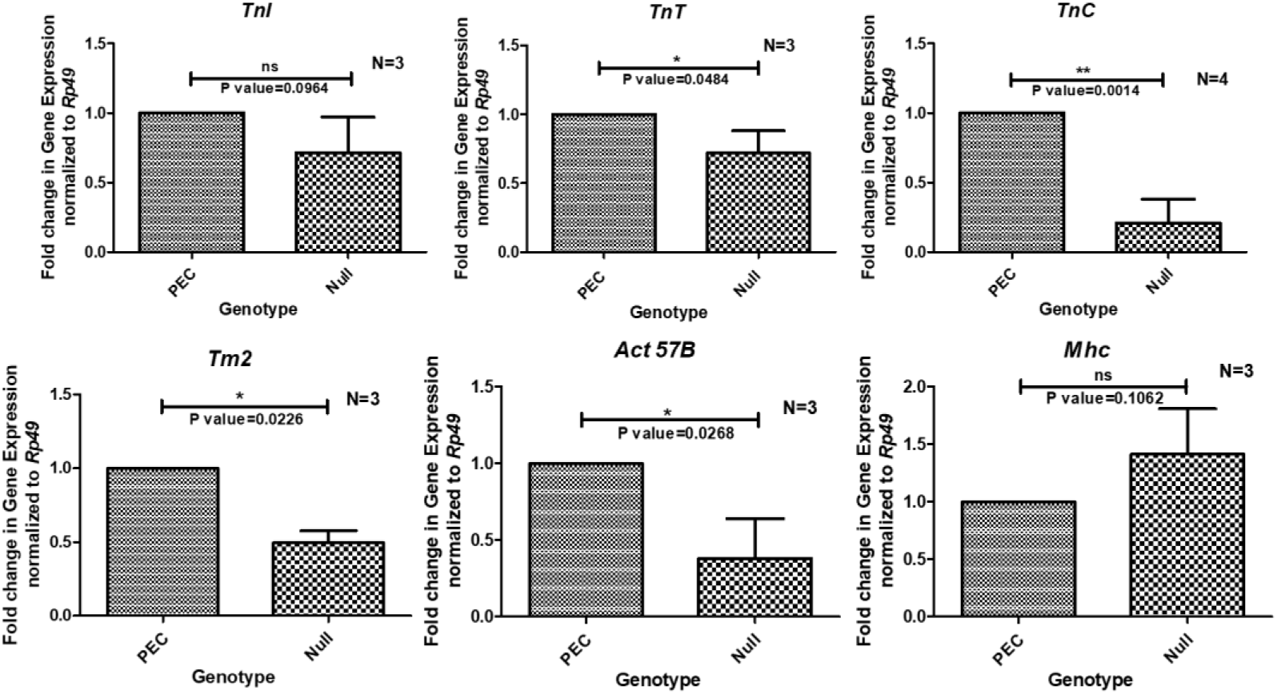
Relative expression (measured by qRT-PCR, using the delta (delta Ct) method) of some major sarcomere protein coding genes in *Mlp60A^null^* L3 larvae. Transcript levels of most of the thin filament proteins we down-regulated.

### Mlp60A-long isoform-specific mutants show drastically compromised flight ability

To determine the function of the *Mlp60A-long* isoform, it was imperative to characterize both the full-length *P-element* insertion allele in the Bloomington line *BS#27970* (renamed *Mlp60A^P-ele^*, to denote the presence of a full length *P{EP}* insertion, and referred to in data panels as, ‘*P-ele*’) and the imprecise hop out allele *UR 2.10.7* (Fig. S6, renamed *Mlp60A^HFDE^*, short for ‘Homozygous Flight Defective Eclosing’ ones, and referred to in data panels as ‘*HFDE*’). We confirmed the insertion of the ~8kb full length *P{EP}* element within the 3^rd^ exon of the *Mlp60A* locus, in *Mlp60A^P-ele^* homozygous flies (Fig. S4). As shown in Fig. 7A, PCR, performed using genomic DNA, isolated from *Mlp60A^HFDE^* homozygous individuals, as the template, and primers covering the insertion site, produced an amplicon of greater size (~1.3kb higher) than those obtained from both the wild type and the *PEC* homozygous flies. The proportion of homozygous *Mlp60A^HFDE^* flies, in the *Mlp60A^HFDE^//CyO-GFP* balanced stocks, was found to be significantly lesser than that expected according to the classical Mendelian ratio for a monohybrid cross (which is followed by the positive control) (Fig. 7C). However, when complementation tests were performed between the different alleles, flies with both *Mlp60A^HFDE^//Mlp60A^null^* and *Mlp60A^HFDE^//Df* genotypes were obtained as per the expected (shown by the positive control) Mendelian ratio (Fig. 7C). Thus, the *Mlp60A^HFDE^* allele can complement both the *Mlp60A^null^* allele as well as a genomic deficiency covering the *Mlp60A* region. This shows that the developmental lethality observed upon *Mlp60A^HFDE^* homozygosity cannot be attributed specifically to this allele. This lethality could, probably, result from the homozygosity of a second site mutation on the *Mlp60A^HFDE^* bearing chromosome, which could have been generated during the hop out. The *Mlp60A^P-ele^* allele was already known to be non-lethal, as stable lines of homozygous *Mlp60A^P-ele^* flies have been maintained by not only in our laboratory, but also at the Bloomington *Drosophila* Stock Centre (BS#27970).

**Figure 7:**
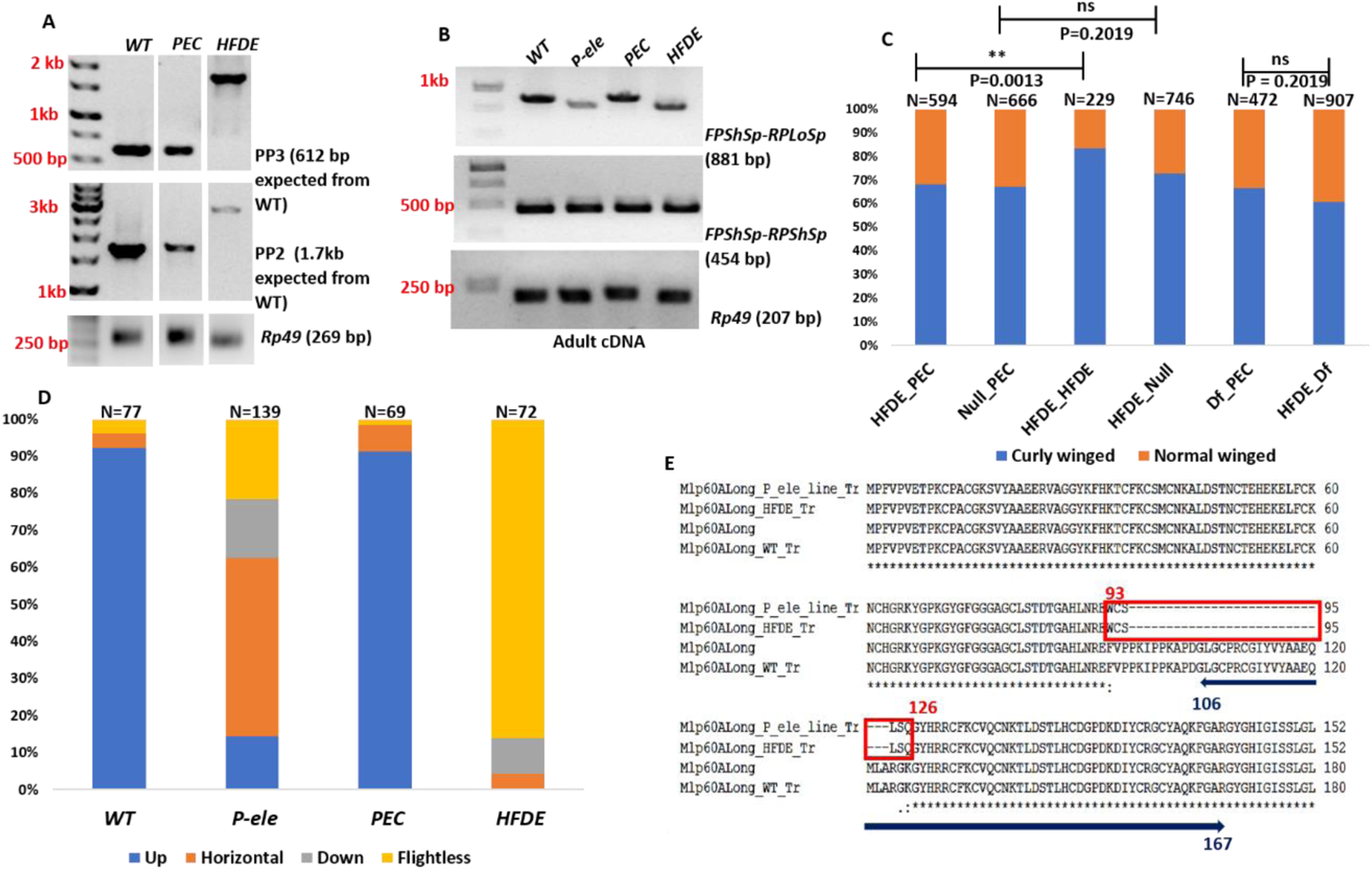
Phenotypic and molecular characterization of Mlp60A-long isoform-specific alleles-*Mlp60A^P-ele^* and *Mlp60A^HFDE^*. (A) Shows the presence of around 1.3 kb remnant of the original *P-element* insertion, in the *Mlp60A^HFDE^* allele. *PP3* refers to PCR products of primers FP3 (intron 2-3) and RP3 (exon 5). *PP2* refers to PCR products of primers FP2 (intron 1-2) and RP2 (exon 7). (B) Shows the analysis of *Mlp60A-long* spliced transcript from homozygous *Mlp60A^P-ele^* and homozygous *Mlp60A^HFDE^* mutants. (C) Shows results of complementation tests between the *Mlp60A^HFDE^* allele and either the *Mlp60A^null^* allele or a *Mlp60A* genomic deficiency chromosome: *Df(2R)BSC356*. Each bar shows the percentages of curly winged and normal winged flies (Y-axis) eclosed in genetic crosses between *CyO-GFP* balanced males/females of respective alleles (X-axis). Homozygous *Mlp60A^HFDE^* flies survive in significantly lesser percentage than the *Mlp60A^HFDE^//PEC* flies (CSV=10.340, df=1), however the survival of *Mlp60A^HFDE^//Mlp60A^null^* flies and *Mlp60A^HFDE^//Df* flies was not significantly different from the respective controls (CSV=0.2019, df=1, in both cases) (D) Shows the flight ability of flies of respective genotypes (all homozygotes). Y-axis shows the percentage of flies and X-axis shows the genotype of flies. (E) Shows the alignment of the translated Mlp60A-long sequences encoded by the *Mlp60A^P-ele^*, *Mlp60A^HFDE^* and *wild-type* alleles, along with the database sequence as the reference. The red box shows the sequence of residues of the long isoform which is different between the wild-type and the mutant long isoform proteins. Residues 93^rd^ to 126^th^ of the wild-type protein are absent from the *Mlp60A^P-ele^* and *Mlp60A^HFDE^* encoded long isoforms. The mutant proteins instead contain unique sequence of six residues-WCSLSQ. The navy-blue bar labels the sequence of the residues in the wild type long isoform, which encodes the 2^nd^ LIM domain. Thus, the comparison presented in this figure clearly shows that the *Mlp60A^P-ele^* and *Mlp60A^HFDE^* allele encoded mutant long isoform lacks part of the 2^nd^ LIM domain.

Next, the *Mlp60A^P-ele^* homozygous flies and *Mlp60A^HFDE^* homozygous flies were tested for their flight ability (Fig. 7D). Around 37% and 48% of the *Mlp60A^P-ele^* homozygous flies were found to be flight-defective and weak-flighted, respectively (Fig. 7D). The flight defect was completely rescued in the *PEC* homozygous flies, whose flight ability was comparable to that of the wild type flies (Fig. 7C, compare 2^nd^, 3^rd^ and 1^st^ bars). In the *PEC* flies the *Mlp60A* locus is restored to the wild type sequence, due to precise excision of the *P-element* from the original insertion site in the *Mlp60A^P-ele^* allele (Fig. S5). On the other hand, as many as ~95% of the *Mlp60A^HFDE^* homozygous flies were flight defective, and the remaining 5% weak flighted (Fig. 7D).

To determine if the presence of the transposon insertions within these two long isoform mutant alleles affect the transcription from the locus, we performed RT-PCR with primers covering the splice site and the insertion site. As shown in Fig. 7B, when cDNA samples, prepared from both the *Mlp60A^P-ele^* homozygous flies and *Mlp60A^HFDE^* homozygous flies, were used as templates, the amplicons obtained appeared slightly truncated as compared to the ones from the wild type and *PEC* homozygous flies. As expected, the amplicons corresponding to the short isoform transcript, obtained from the *Mlp60A^P-ele^* homozygous flies and *Mlp60A^HFDE^* homozygous flies, were identical in size, to those obtained from the wild type and *PEC* homozygotes (Fig. 7B). The sequences of amplicons corresponding to the long isoform transcripts, encoded by the *Mlp60A^P-ele^* and *Mlp60A^HFDE^* alleles respectively, when aligned (Fig. S11) with each other, were found to be completely identical (GenBank Accession no: MN990116). These transcript sequences were then aligned with the sequence of the corresponding wild type transcript, to determine the extent of the truncation (seen in Fig. 7B). Fig. S12 depicts the analysis of the region flanking the original insertion site. As shown by Fig. S12 A, both of the transcripts contain a unique stretch of 17 bases, which is not represented in the wild type transcript sequence (shown by purple ovals), and both are devoid of a stretch of sequence, which is present in the wild type transcript (shown by orange rectangles). A further analysis (Fig. S12B) revealed that these transcripts resulted from an alternative splicing event, in which the usual splice acceptor is avoided and a novel splice acceptor, preceding the 4^th^ exon is selected instead. However, a normal splice donor is generated resulting in the splicing out of the 3^rd^ exon and the inclusion of a sequence of 17 bases (1726^th^-1742^nd^ base) from intron 3-4, in the mature spliced transcript, thus ensuring that the insertion in the 3^rd^ exon does not lead to a complete disruption of the reading frame (Fig. S12B). Protein sequences from both these truncated transcript sequences when aligned with the sequence of the wild type Mlp60A-long protein (Fig. 7E) led to the removal of a total of 34 amino acid residues (93^rd^ to 126^th^ residues), encoded by the 3^rd^ exon. Of these, 21 residues (106^th^ to 126^th^ residues) belong to the second LIM domain of the longer isoform. Also, 6 unique residues (WCSLSQ), encoded by the extra sequence of bases in the mature transcript, are included in the protein sequence. Hence the *P-element* insertion in both these alleles leads to the expression of a mutant Mlp60A-long isoform, which has a truncated and modified second LIM domain. Overall, the flight phenotype of the *Mlp60A^P-ele^* homozygous flies and its subsequent rescue in the homozygous *PEC* flies, strongly suggests that the wild type Mlp60A-long isoform with all intact LIM domains is essential for normal flight.

### *Mlp60A^P-ele^* homozygotes show normal IFM myofibrillar structure, but distorted sarcomere length

To test if the Mlp60A-long isoform was required for maintenance of flight in an age-dependent manner, we tested the flight ability of *Mlp60A^P-ele^* homozygotes at both 3-5 days and 10-12 days of age. As shown by Fig. 8A, there was an increase in the percentage of flight defective *Mlp60A^P-ele^* homozygous flies, from 34% to 48%, from 3-5 days to 10-12 days of age, whereas the flight ability of the control flies remained almost the same. The longitudinal sections of the IFMs from some of the 10-12 days old, flight defective, *Mlp60A^P-ele^* homozygous flies were processed for visualization of their myofibrillar structure. As shown in Fig. 8B, there was no defect in the myofibrillar architecture of the IFMs from *Mlp60A^P-ele^* homozygous flies. However, there was a significant increase in sarcomere length (1.7%, calculated by difference of means) of the IFMs in these flies, as compared to the *PEC* homozygous flies (Fig. 8C). In the *Mlp60A^P-ele^* homozygous flies, all the sarcomeres quantified were greater than 3 µm (Note the distribution of points in Fig. 8C). This data suggests that the long isoform is necessary for the achievement of normal sarcomere length, during the maturation phase of IFM myofibrils, which occurs after eclosion [48].

**Figure 8:**
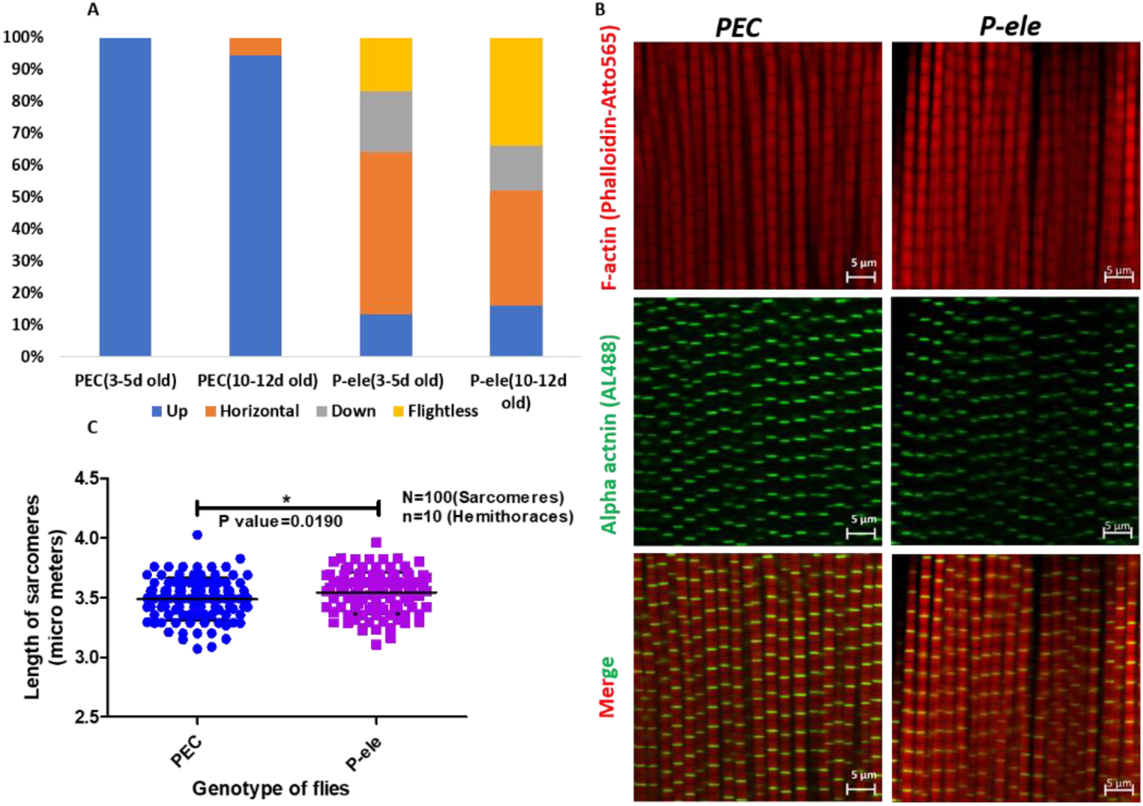
Myofibrillar defects in *Mlp60A^P-ele^* homozygous mutants. (A) Shows the flight profile of *Mlp60A^P-ele^* homozygotes. (B) Representative images of IFM myofibrils from homozygous *Mlp60A^P-ele^* and *PEC* flies, stained for F-actin filaments (red channel) and α-actinin (green channel) *Mlp60A^P-ele^* homozygotes show normally aligned myofibrils, but the sarcomere length of these are slightly longer than the corresponding control, as quantified in (C). Scalebar reads 5 µm.

## Discussion

The Muscle LIM protein has long been regarded as a differentiation factor, which promotes the differentiation of myoblasts into myocytes in culture [31–32, 49]. However, it’s role during the development of embryonic muscles, which are the first differentiated muscle structures *in-vivo* [50], has not been studied in detail. Our results show that this protein is necessary for the development of the larval body wall muscles of *Drosophila*, which are a type of embryonic muscles [51]. Since the locomotion defect and body wall muscle defects in the *Mlp60A^null^* mutants were majorly seen at the L3 stage and not at the earlier stages (data not shown), it is very likely that the *Mlp60A-short* isoform could be required for the maintenance of the larval body wall muscles. It is known that the body wall muscles, after initial development in the embryo, increase drastically in size (~25 to 40-fold increase in area) by hypertrophic growth, over the next 5 days, as the development proceeds from L1 to L3 stage [52]. The myofibrillar-contractile proteins form a major portion of the dry weight of muscles, and an increase in muscle size is majorly due to increase in the contractile protein content [53]. We found that the expression of a few major sarcomeric structural proteins, particularly thin filament proteins, are significantly reduced in *Mlp60A^null^* homozygotes (Fig. 6). Therefore, it is likely that the complete absence of Mlp60A, a thin filament protein [49, 54–55] and the resultant reduction in the amounts of other major thin filament proteins, such as Actin(57B), Troponin I, Troponin T, Troponin C and Tropomyosin 2, leads to the reduction of muscle thickness and muscle loss, seen in the Mlp60A null mutants. However, Myosin heavy chain expression was not affected in the Mlp60A null mutants, unlike that reported by Rashid *et al*., upon *in vitro* depletion of the MLP [56]. It is important to note here that the phenotype shown by the *Mlp60A^null^* homozygotes cannot result from the partial *CG3209* disruption, which the *Mlp60A^null^* allele bears, since the *CG3209* loss of function mutants, characterized by Yan *et al*., did not show any post-L1 lethality or L3 stage locomotion defects [57]. Therefore, it can be concluded that the Mlp60A-short isoform is essential for the development of the larval body wall muscles. The other MLP ortholog Mlp84B, on the other hand, does not appear to play any role in the development of the larval body wall muscles since the Mlp84B null mutants do not show any phenotype until the onset of pupal development [58].

The expression profile of *Mlp60A* isoforms (Fig. 1C-E) mimics that of several other sarcomeric proteins, which have IFM or IFM-TDT specific isoforms, that are selectively expressed in the IFMs during their remodelling from the larval muscles, with the embryonic/pupal isoforms being down-regulated [22, 25, 27–30]. Such an expression profile suggests the presence of functional specialization among these isoforms, as has been found to be true for other sarcomeric proteins [11, 24, 30, 59–60]. Our results show that the Mlp60A-long isoform, with one truncated LIM domain, is necessary for achieving normal sarcomere length in the IFMs. Longer than usual sarcomere length is known to result in sub-optimal force production since it reduces the effective overlap between the actin and myosin filaments during muscle contraction, as explained by the sliding filament model [61]. This could be the most likely reason for the reduced flight ability of the *Mlp60A^P-ele^* homozygous flies. This result is consistent with the function of other sarcomeric proteins such as Myosin heavy chain (Mhc) and Actin 88F (Act88F), in the maintenance of sarcomere length in adult IFMs [62]. Interestingly, however, the sarcomere length in the IFMs was significantly reduced in the *Mlp60A* knockdown flies. It is known that shorter sarcomeres lead to production of significantly lesser force (61). The IFM knockdown phenotype being very drastic in nature, the reduced sarcomere length could be a compensatory response of the system, to prevent further muscle damage, by reducing the force produced with each contraction cycle.

Several studies conducted previously have shed light on the functional non-redundancy of the different isoforms of myofibrillar proteins, in Drosophila. Several dominant flightless, point mutants of the *Act88F* gene, which encodes for the IFM specific actin isoform-Act88F, have been isolated [63–64]. Further, it has been shown that the IFM specific isoform-Act88F can functionally compensate for the loss of the TDT specific isoform-Act79B, but the converse is not true [65]. The IFM-TDT specific TnI null mutant-*hdp^3^* shows loss of flight and jumping ability, with a drastic hypercontraction phenotype in the IFMs [22, 30]. Also, Nongthomba *et al*. have shown that the IFM-TDT specific TnT null mutant-*up^1^*, shows loss of only flight and jump ability, and structural defects in IFM and TDT, but normal larval crawling and adult walking ability (25). Studies conducted on the Drosophila MHC locus have shown that the different, alternatively spliced MHC isoforms, possess differences in their relay or converter domains [66–67] or S2 hinge region [68–69] or N-terminus region [70–71]. These functional domains of the IFM specific MHC isoform, cannot be functionally compensated by the respective domains of the embryonic MHC isoforms. These studies show the necessity of expressing the correct isoforms of different myofibrillar proteins in the IFMs for normal physiological function. Our results show that ectopic expression of the short isoform in the IFMs, can weakly compensate for the complete absence of the long isoform in the IFMs, with regards to flight ability. This feeble functional compensation could result either from sub-normal expression levels of the short isoform or from an inability of the short isoform to compensate for the long isoform function. However, no such functional compensation was seen in the homozygous *Mlp60A^P-ele^* mutants, which express a normal short isoform, suggesting that it could be solely due to the ectopic expression of the short isoform and that the longer isoform is necessary for normal IFM function. On the other hand, a simultaneous knockdown of both these isoforms leads to an even more drastic flight defect and IFM defects, suggesting that the longer isoform is the major one governing normal IFM function and hence, flight ability. In the absence of the long isoform, the short isoform can provide for only a weak flight ability. In vertebrates, Vafiadaki *et al*., have reported an alternate isoform of MLP, designated as MLP-b (to differentiate it from the known isoform, renamed MLP-a). This isoform is generated as a result of splicing out of exons 3 and 4 from the primary *CSRP3* transcript [41]. MLP-b levels were upregulated in limb girdle muscular dystrophy (LGMD2A), Duchenne muscular dystrophy (DMD) and Dermatomyositis (DM) patients and the MLP-b/MLP-a ratio was found to be altered in LGMD2A and DMD patients [41]. These results show that the deregulation of MLP alternative splicing can contribute to the pathogenesis of these diseases. Also, deregulated splicing has been implicated in diseases like Myotonic Dystrophy-type1 (MD-1), fascioscapulohumeral dystrophy (FSHD), DCM and HCM [72–75]. Most notably, it is known that the loss of function of human Muscleblind proteins, due to their binding to *CUG*-repeat bearing RNA foci of the *DMPK* transcripts in the nucleus, leads to altered splicing of several primary transcripts such as those of cardiac Troponin T (cTnT) and the Insulin Receptor (IR) [75–78]. Taken together, these results point towards the importance of identifying splicing factor(s) responsible for the alternative splicing of *CSRP3* primary transcript, to gain a deeper understanding of the pathogenesis of diseases in which MLP isoform levels are altered. A future study of the players involved in *Mlp60A* alternative splicing in *Drosophila* IFMs can greatly aid in dissecting this process.

*CSRP3* gene is usually included on the list of genes for analysis of mutations in HCM patients, as several *CSRP3* variants have been reported, which have been linked to this disease [35–38, 79–80]. However, till date only one variant, the p.C58G variant, in the 1^st^ LIM domain of MLP has been validated to be ‘likely pathogenic’, through *in vivo* studies [38]. Our results show that the loss of a part of the 2^nd^ LIM domain of MLP affects the sarcomere length which is detrimental to the physiological function (in our case, flight), thus providing additional *in-vivo* evidence that MLP variants may play an important role in the development of HCM phenotype, which needs to be studied further.

Additionally, the IFM phenotype of the *Mlp60A* knockdown flies (frayed myofibrils and formation of actin rich aggregates), shows that this locus is involved in the regulation of actin dynamics and thin filament assembly, during IFM development. These results corroborate those of a previous study wherein MLP was shown to promote actin crosslinking and bundling of actin filaments, in C2C12 myoblasts [55]. In conclusion, we have shown that the two isoforms of Mlp60A in Drosophila not only show spatio-temporally regulated expression but are also functionally non-redundant.

## Materials and Methods

### Fly lines used in the study

The fly lines used in this study were either procured from the Bloomington Drosophila Research Center (Bloomington University, Indiana) or the Vienna Drosophila Research Center (Vienna BioCenter, Vienna), or were generated in the lab. The following Stocks were used:

i. Mlp60A RNAi line1: BS#29381 (y[1] v[1]; P{y[+t7.7] v[+t1.8]=TRiP.JF03313}attP2).
ii. Mlp60A RNAi line2: VDRC#23511 (w^1118; P{GD13576}v23511^). Fly lines (i) and (ii) were used to achieve the RNAi-mediated knockdown of *Mlp60A* by the *UAS/Gal4* system [43].
iii. *Drosophila* line containing a source of the genetically engineered *P-element* transposase: “delta 2-3 transposase”: BS#4368; (*y*[1] *w*[1]*; Ki*[1] *P{ry[+t7.2]=Delta2-3}99B*).
iv. Mlp60A P-element insertion line: BS#27970; (y[1] w[*]; P{w[+mC]=EP}Mlp60A[G7762]) [81]. Fly lines (iii) and (iv) were used to perform a hop out mutagenesis screen, to generate null or isoform-specific *Mlp60A* mutants.
v. Dmef2-Gal4 driver line: BS#27390; (y[1] w[*]; P{w[+mC]=GAL4-mef2.R}3) [82]
vi. elav-Gal4 driver line: BS#458; (P{w[+mW.hs]=GawB}elav[C155]) [83].
vii. *UH3-Gal4* driver line: generated in the lab [45]. Fly lines (v-vii) were used as the *Gal4* driver lines to achieve tissue-specific knockdown or overexpression of the *Mlp60A*.
viii. A deficiency line of *Mlp60A* region: BS#24380; (*w1118; Df(2R)BSC356/SM6a*). Apart from these lines, the various balancer chromosomes, wherever required, were used as per the standard methodology of genetic crosses for *Drosophila* [84].

### Maintenance and culturing of flies

The flies were cultured using the standard *Drosophila* medium (Cornmeal-Agar-Sucrose-yeast) at 25^0^C. Crosses for tissue-specific knockdown or over-expression using the *Gal4/UAS* system were carried out at 29^0^C. The larvae wherever required, were collected and reared on a larval collection medium with the following composition: 2.5% ethanol, 1.5% Glacial Acetic Acid, 1.5% agar, 2.5% sucrose; with a thick yeast paste as the major food source.

### Genetic complementation tests and rescue

The complementation tests were carried out by crossing together the *CyO-GFP* balanced fly stocks of the respective alleles (or the deficiency chromosome). The percentages of trans-heterozygous (normal winged) and curly winged flies obtained in the resulting progeny was compared with those obtained in the respective control cross, in which one of the test stocks was crossed to the *PEC//CyO-GFP* fly stock, to yield normal winged and curly winged flies according to Mendelian ratios.

The rescue experiment was performed using a similar strategy. The flies of *Mlp60A^null^//CyO-GFP; UAS-Mlp60A-short* genotype were crossed with flies of either genotypes: *Mlp60A^null^//CyO-GFP* (negative control set), or *PEC//CyO-GFP* (positive control set), or*UH3-Gal4; Mlp60A^null^//CyO-GFP* (test set). The percentages of normal and curly winged flies obtained in each set were then compared to assess the extent of rescue.

### Genetic crosses to generate *P-element* hop outs

Through the appropriate genetic crosses, the *P-element* insertion bearing chromosome and the chromosome carrying the transposase gene were brought together. Flies of this genotype were mated with flies carrying the 2^nd^ chromosome balancer: *Tft//CyO*. From the progeny, each white eyed male fly (hop out) was mated with *Tft//CyO* virgin flies in a separate vial, to obtain stable lines. The progenies from such crosses (stable lines) were then subjected to ‘selfing’ and screened for phenotypes.

### Flight test

Flies were tested for their flight ability using the ‘Sparrow box’ method as described previously [62]. Each fly was tested 2-3 times and results were presented as percentages of flies showing each type of flight ability-Up, Horizontal, Down or Flightless, as per their flight in the Sparrow box. Unless otherwise mentioned, all flight tests were performed with 3-5 day old flies.

### Test of larval locomotion

Third instar larvae were tested for their crawling ability on 1% agar plate. The animals were transferred onto a moist 1% agar plate (90 mm dish) and allowed to acclimatize for about 30s. Following this, they were allowed to crawl for 42s, during which time videos were captured using a digital camera (8 Mega Pixel). The recorded videos were then analysed using the “MtrackJ” plugin with ImageJ v1.52k (https://imagej.nih.gov/ij/).

### Visualization of IFMs through polarized light microscopy

The fly hemi-thoraces were processed for visualization of IFM fascicular structure by polarized light microscopy as described previously [85–86]. Following preparation, the hemi-thoraces were observed using an Olympus SZX12 microscope with a polarizer and analyser attachment. Images were captured using a Leica DFC300 FX camera.

### Dissection and visualization of larval body wall muscles by confocal microscopy

Third instar larvae were immobilized by placing them on ice in a cavity block for 30 minutes. These were then dissected by placing them in 1X PBS on a Silguard plate. Following this, the larvae were fixed by using 70% alcohol for 30 min washed with 1X PBS, 3 times, on a rocker and then stained with 1:40 diluted Phalloidin-TRITC (Sigma) for a period of 1 hr at room temperature. Then the larvae were washed again using 1X PBS and then mounted on a slide using 1:1 mixture of Glycerol and Vectashield (https://vectorlabs.com/vectashield-mounting-medium.html) as the mounting agent. The cover slips were then sealed with transparent nail paint and then stored at 4^0^C. These samples were then visualized under a confocal microscope (either Zeiss LSM 510 META or Leica SP8 confocal imaging system).

### Visualization of IFM structure by immunostaining and confocal microscopy

Dissections and sample preparation for confocal microscopy to visualize the transverse (cross) sections and longitudinal sections of IFMs were performed as described previously [87]. The Z-discs were marked in the longitudinal sections by using mouse anti *α*-actinin primary antibody (DSHB Hybridoma, Product 47-18-9). The muscle membrane was marked using mouse anti *β-*PS-Integrin primary antibody (DSHB Hybridoma, Product CF.6G11). Both the primary antibodies were used with 1:100 dilution. The secondary antibody used in both cases was anti-mouse AL488 (Invitrogen), at 1:250 dilution. Primary Anti-Mlp60A (raised in rabbit, raised in Lab) antibody was used with 1:1000 dilution. Samples were visualized using either Zeiss LSM 510 META or Leica SP8 confocal imaging system.

### Assessment of developmental lethality

To assess the effective lethal stage of mutants, eggs were collected on a 2.5% sucrose, 1.5% agar medium in 60 mm petri plates. Following the collection of eggs, they were allowed to hatch and the L1 larvae were transferred to an egg collection medium with an additional 1.5% glacial acetic acid and 2.5% ethanol. Then their development was monitored to check up to which stage each individual survived. The lethality was calculated as per the percentage of total fertilized eggs or L1 larvae initially collected, that died in the subsequent larval or pupal developmental stages [88].

### Genomic DNA isolation

Genomic DNA was isolated from either whole flies or whole larvae using the Qiagen ‘DNeasy Blood and Tissue’ DNA isolation kit (Catalogue no:69504). The isolated genomic DNA was quantified by using NANODROP 1000 or NANODROP Lite spectrophotometers by ThermoFisher Scientific.

### RNA isolation and cDNA preparation

Either whole animal (adults, L1 larvae, L3 larvae or pupae of different stages) or tissues (adult head, thorax or abdomen or IFMs) were collected for RNA isolation using the TRI reagent (Sigma) based protocol. The isolated RNA was quantified by using NANODROP 1000 or NANODROP Lite Spectrophotometers by ThermoFisher Scientific. The integrity of the isolated RNA was assessed by running 1 μL of isolated RNA on 1% agarose gel. Reverse transcription reaction was carried out using the first-strand cDNA synthesis kit by ThermoFisher Scientific.

### Qualitative Polymerase Chain Reactions

PCRs were carried out using 2X PCR Master mix (Taq Polymerase) by ThermoFisher Scientific (Catalogue No: K1072). High fidelity PCRs, wherever required, were carried out by using ThermoFisher Scientific Phusion DNA polymerase (Catalogue No: F530S). The products resulting from PCR were resolved on agarose gels of different compositions (0.8%, 1% or 2%) and imaged.

### Quantitative Polymerase Chain Reactions

qRT-PCRs were carried out to assess relative gene expression in different samples. The reactions and data analyses were carried according to the ‘delta (delta)C_t_’ method as described by Livak and Schmittgen, 2001 [89]. In cases, wherever required, only delta C_t_ was plotted. The reactions were carried out using the ‘SYBR Green Master Mix’ by Bio-Rad or the ‘DyNamo SYBR Green qPCR kit’ by ThermoFisher Scientific (Catalogue No: F-410L).

### Sequencing of DNA fragments

DNA fragments amplified by PCR and sequenced either directly after performing a clean-up with ‘QIA Quick PCR Purification Kit’ (QIAGEN Catalog No: 28104) or sequenced after generating clones (TA-cloned using the ‘ThermoFisher Scientific InsTAclone PCR Cloning Kit’, Catalogue No: K1214). Samples were sequenced at AgriGenome Labs.

### Generation of transgenic *UAS* lines for conditional expression of *Mlp60A-short* isoform

The Mlp60A-short specific CDS was cloned within the pUASt-attB vector following the usual methodology for restriction digestion-based cloning. The *pUASt-attB-Mlp60A-short* plasmid construct was submitted to ‘Fly Facility, NCBS’ (http://www.ccamp.res.in/Fly-facility), for microinjection into embryos carrying *attP2* docking site on the 3rd chromosome for *attP2-attB* mediated site-specific insertion of the constructs. The microinjection and screening for transgenics were performed by the Fly Facility. Two transgenic lines were obtained, one of which was used for the experiments.

### Molecular Cloning

DNA fragments were cloned by restriction digestion-ligation based procedure, for which all enzymes were obtained from ThermoFisher Scientific. *E. coli* DH5α competent cells were prepared by the TSS protocol [90]. The transformation was done using the heat-shock protocol [91]. The colonies were screened by PCR with insert specific primers, following which plasmids were isolated using the ‘GSURE Plasmid Mini’ kit by GCC Biotech (Catalogue No: G4613).

### Raising Polyclonal antibody against Mlp60A smaller isoform

The polyclonal antibody was raised against the smaller Mlp60A isoform of 10 kDa size by cloning the 200 bp transcript into the pET15b protein expression vector after confirming the sequence (Macrogen, South Korea) for any mutation. Then, the protein was expressed in the *E. coli* (BL21 (DE3) endo strain upon induction using 0.4 mM isopropyl B-D-1-thiogalactopyranoside (IPTG) for 6 hrs at 25^0^C and injecting the expressed product into a rabbit to raise antibody.

### Protein extraction and Western blot

IFMs were dissected from bisected flies preserved in 70% alcohol and homogenized in 1x Buffer (0.1 M NaCl, 10 mM potassium phosphate pH 7.0, 2 mM EGTA, 2 mM MgCl_2_, 1 mM DTT, 1 mM PMSF and 0.5% Triton-X). The IFM lysate was spun down to obtain a protein pellet which was further washed with the same 1X buffer but without Triton-X and then boiled in SDS-sample buffer (0.0625 M Tris-Cl pH 6.8, 2% SDS,10% glycerol, 5% 2-mercaptoethanol and 5 µg bromophenol blue) for 4 minutes at 95°C. Samples were then resolved in a 12% PAGE gel in mini electrophoresis unit (Amersham) at 100V. The protein was then transferred from the gel to a PVDF membrane (Immobilon-P, Millipore) in transfer buffer (20% methanol, 25 mM Tris-base and 150 mM glycine). The membrane was blocked with 8% milk solution in Tris buffer saline (TBS, pH 7.4) for 1 hr and then probed with the primary antibody at prescribed dilution overnight at 4°C. Anti-Mlp60A (raised in rabbit, raised in Lab) was used with 1:1000 dilution. After washing three times with TBS, the membrane was incubated with the HRP-conjugated secondary antibody (anti-rabbit 1:1000, Bangalore Genei, Bangalore) for 3 hrs at room temperature. The membrane was then washed three times (15 minutes each) with TBST (TBS with 0.05% Tween 20) and 5 minutes with 0.5 M NaCl. Bands were detected by using the enhanced chemiluminescence (ECL) method (Supersignal WestPico Chemiluminescent substrate, Pierce).

### Primers used in this study

Supplementary Table S1 lists the primers that were used in this study, along with the respective annealing temperatures for PCR and the purpose for which each was used.

### Quantifications and statistical analyses

Phenotypic parameters were quantified using ImageJ v1.52k (for IFM fascicular width, total fascicular area and sarcomere length quantification) or the LSM Image browser (for larval body wall muscle width measurement). The resulting data were plotted in GraphPad Prism v5.00 and the statistical significance was determined by using unpaired Student’s t-test. Statistical significance for qPCR data was carried out using paired Students t-test. The chi-squared test was carried out to determine statistical significance in data sets with categorical data.

## Acknowledgements

We thank the Bioimaging facility and Animal facility at the Indian Institute of Science (IISc) for their aid. We also thank the University Grants Commission, Ministry of Human Resource Development (MHRD), Govt. of India, for research fellowship (Sr No: 2121330889, Ref No: 22/12/2013(ii)EU-V). This worked was supported by SERB, Department of Science and Technology (DST), Govt. of India, grant (EMR/2016/004563) to UN. We also acknowledge the Indian Institute of Science (IISc), the Department of Science and Technology (DST), Govt. of India, (DST FIST, Ref. No. SR/FST/LSII-036/2014), the University Grant Commission (UGC-SAP to MRDG: Ref. No. F.4-13/2018/DRS-III (SAP-II) and the Department of Biotechnology (DBT), Govt. of India, (DBT-IISC Partnership Program Phase-II (BT/PR27952-INF/22/212/2018) for infrastructure and financial assistance. We also would like to acknowledge anonymous reviewers for their insightful comments and suggestions which allowed us to improve the manuscript.

## Author Contributions

- MJ along with SRR and UN conceived and performed the experiments pertaining to Mlp60A polyclonal antibody generation and the associated Western blotting and Immunostaining.
- RW and UN conceived and planned all the other experiments being reported in the manuscript, and these were performed exclusively by RW.
- Paper was written by RW and UN, and approved by all the authors before submission.

## Supplementary Data

**Figure S1:**
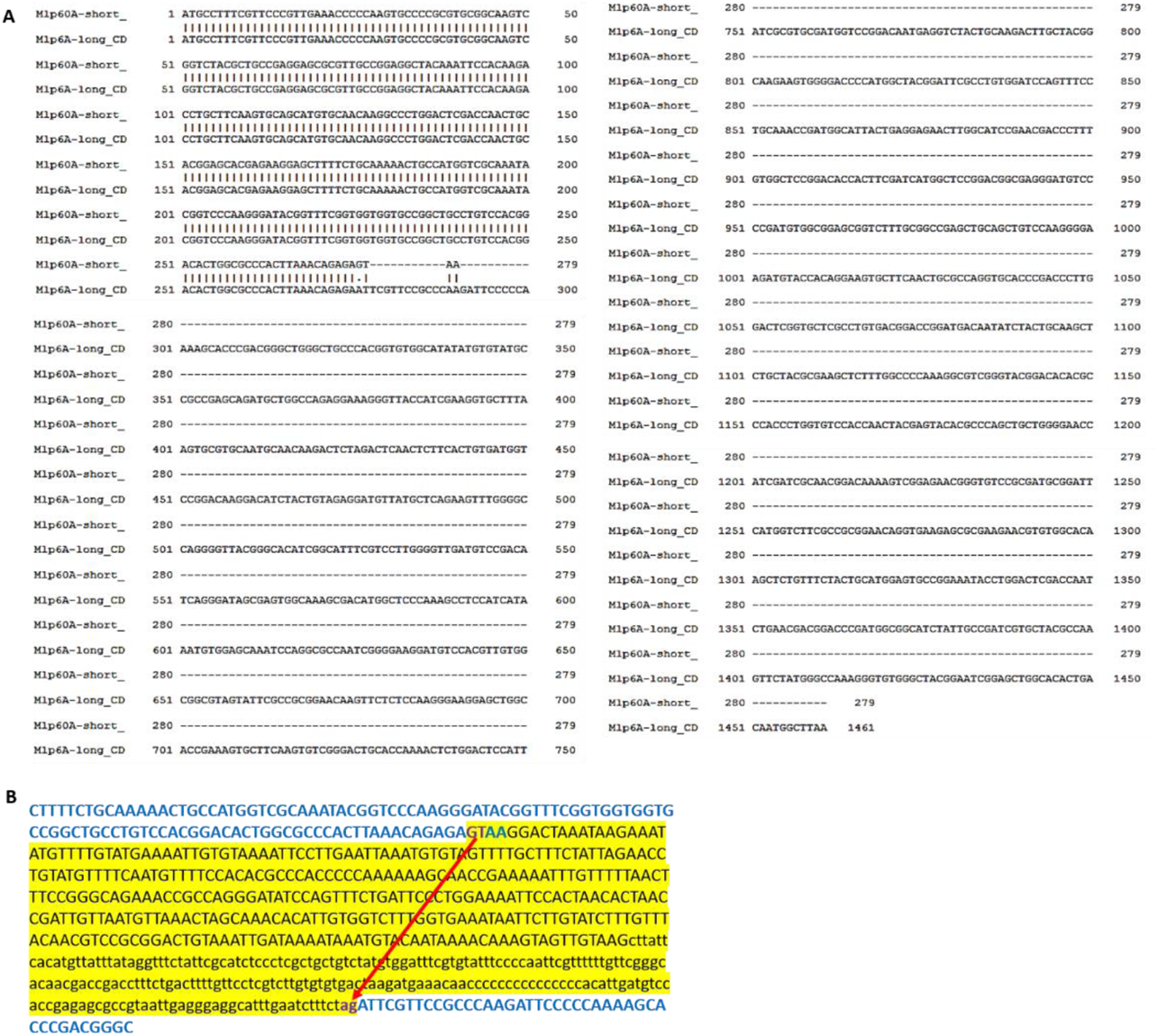
Bioinformatic alignment of *Mlp60A-short* and *Mlp60A-long* Coding Sequences. (A) Shows the complete alignment of both the *Mlp60A* isoforms. (B) Shows a part of the *Mlp60A* genomic sequence, depicting the alternative splicing event which leads to expression of the long isoform. The sequences of the 2^nd^ and 3^rd^ exons have been coloured blue. The short specific 3’UTR sequence has been coloured black and highlighted in yellow colour. The splice donor and acceptor base combinations have been coloured purple. The red arrow marks the splice donor (GT) and acceptor (ag) sites.

**Figure S2:**
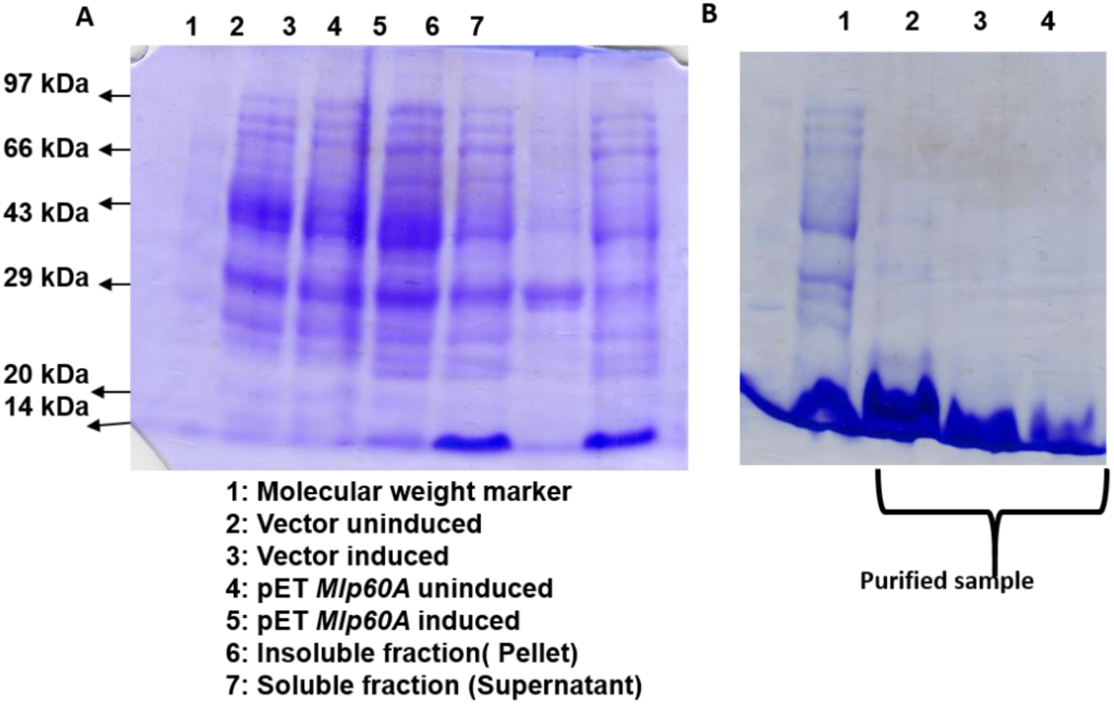
Expression and purification of Mlp60A-short isoform for generation of polyclonal antibodies. (A) Shows the successful induction of protein, as achieved in *E. coli BL21* strain transformed with *pET15b* vector carrying the *Mlp60A-short* CDS insert. The induction was brought about by exposing the cells to 0.4 mM IPTG for 6h at 25 ^0^C. (B) Shows the purified protein isolated by using His-tag purification kit.

**Figure S3:**
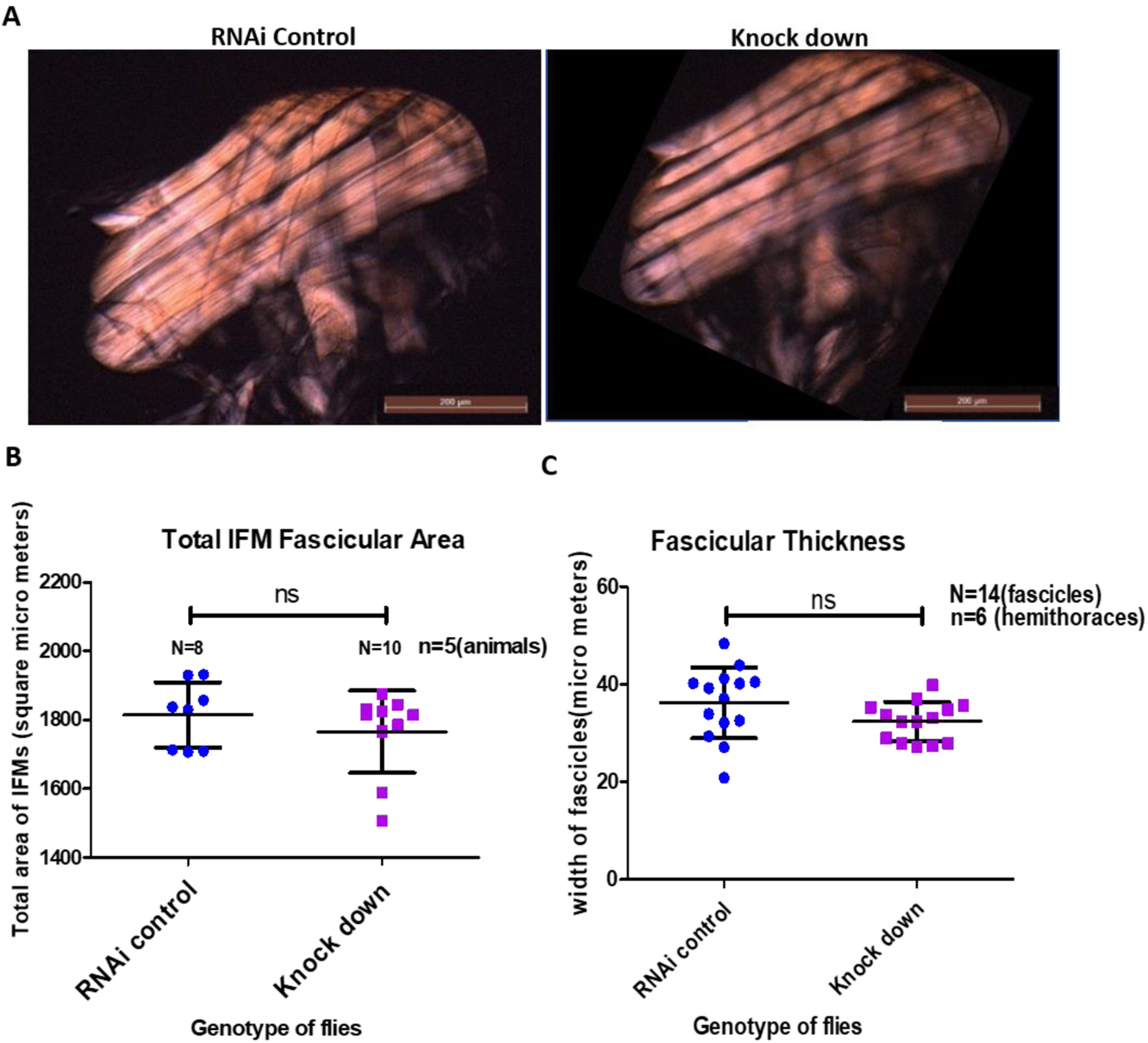
Analysis of IFM fascicular structure in *Mlp60A* knockdown flies. (A) Polarized light images of IFMs from RNAi control and knockdown flies. The scale bar reads 200 µm. (B) Quantification of total IFM fascicular area and (C) IFM fascicular thickness, in *Mlp60A* knock down flies. Genotypes-RNAi Control: *UAS-Mlp60A-RNAi_1(III)*, Knock down: *Dmef2-Gal4(III)::UAS-Mlp60A-RNAi_1(III)*.

**Figure S4:**
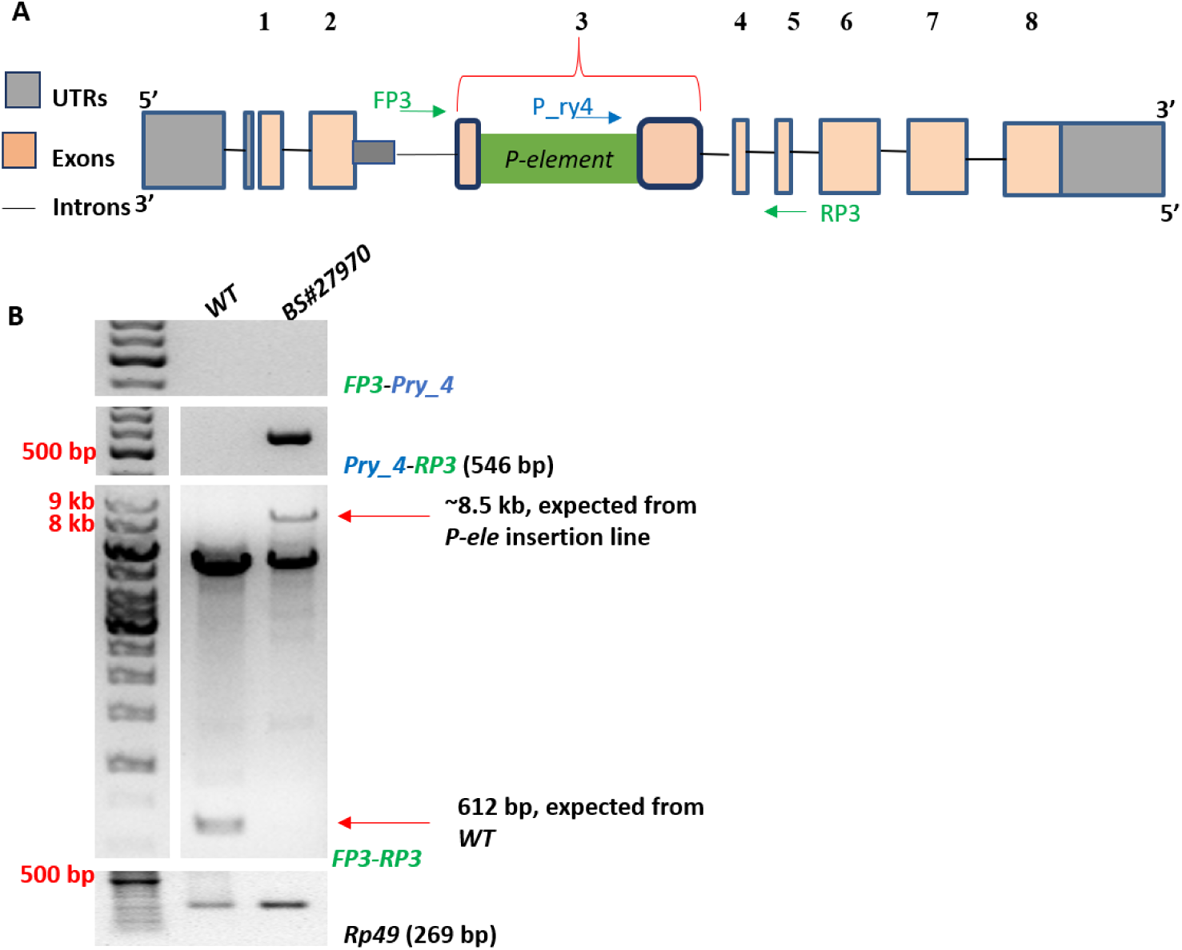
Verification of *P-element* insertion in BS#27970 *Drosophila* line by PCR. (A) Line diagram showing the insertion. (B) PCRs to confirm the position and orientation.

**Figure S5:**
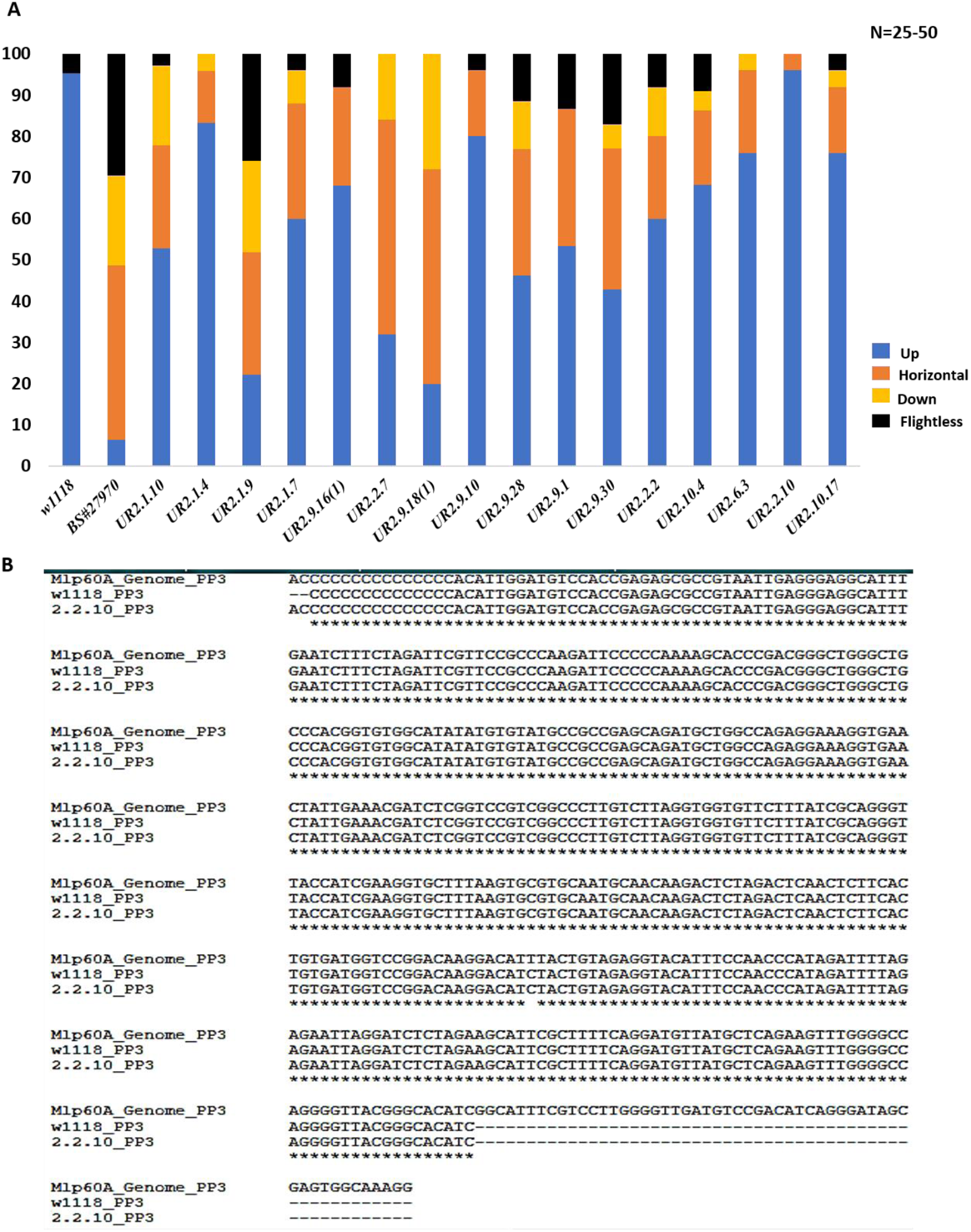
Identification of a Precise Excision Allele. (A) Shows the flight ability of flies homozygous for different candidate alleles. Y-axis shows the percentage of flies, X-axis shows the genotype of flies. (B) Shows the alignment of *FP3-RP3* amplicons (see Materials and Methods for primer positions) from homozygous *UR 2.2.10* flies and wild type flies, along with the database sequence as a reference sequence.

**Figure S6:**
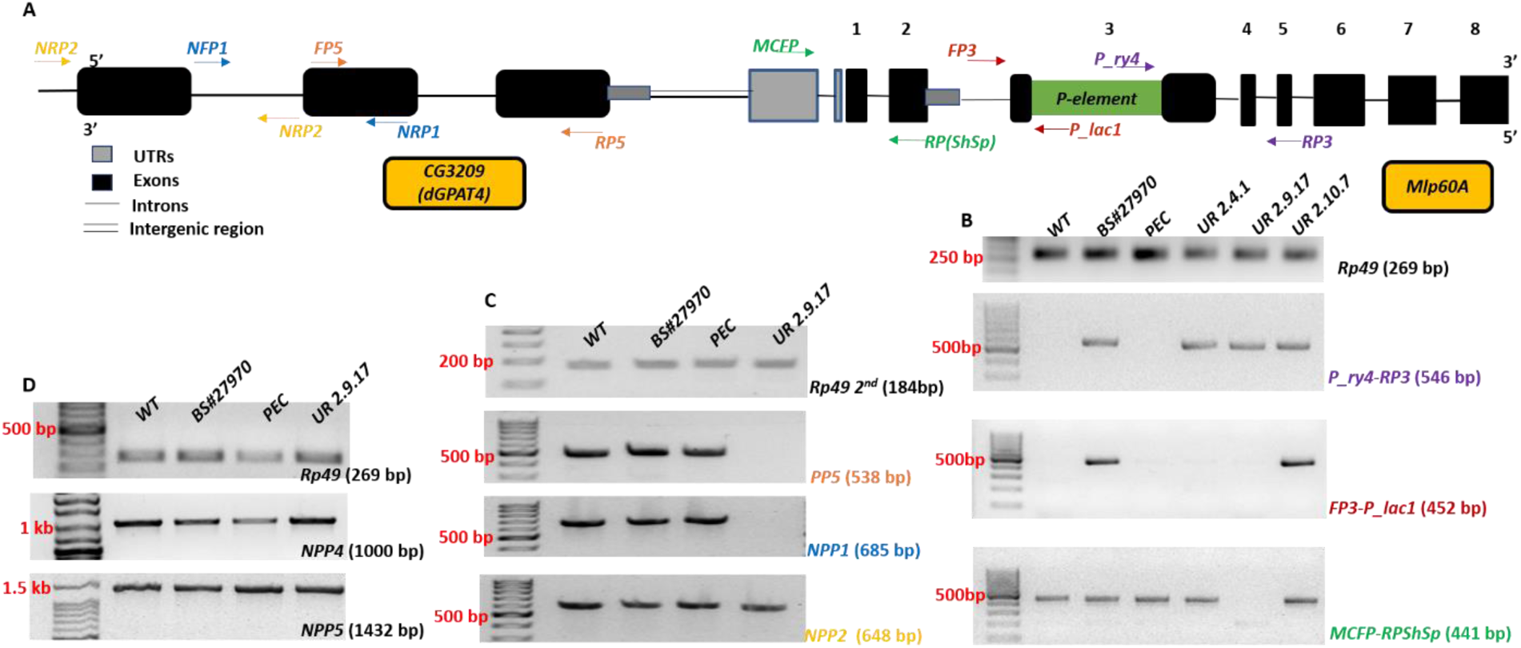
Genomic characterization of homozygous lethal mutant alleles recovered in *P-element* hop out mutagenesis screen. (A) Shows the positions of the different primer pairs used for the analysis, along the Mlp60A locus and the upstream locus. (B-C) Show the PCR products obtained with respective primer pairs from different samples. Each primer pair and its corresponding PCR product have been represented in the same colour. (D) Shows results of PCRs with primer pairs designed further upstream, in the CG3209 region (for positions see Materials and Methods).

**Figure S7:**
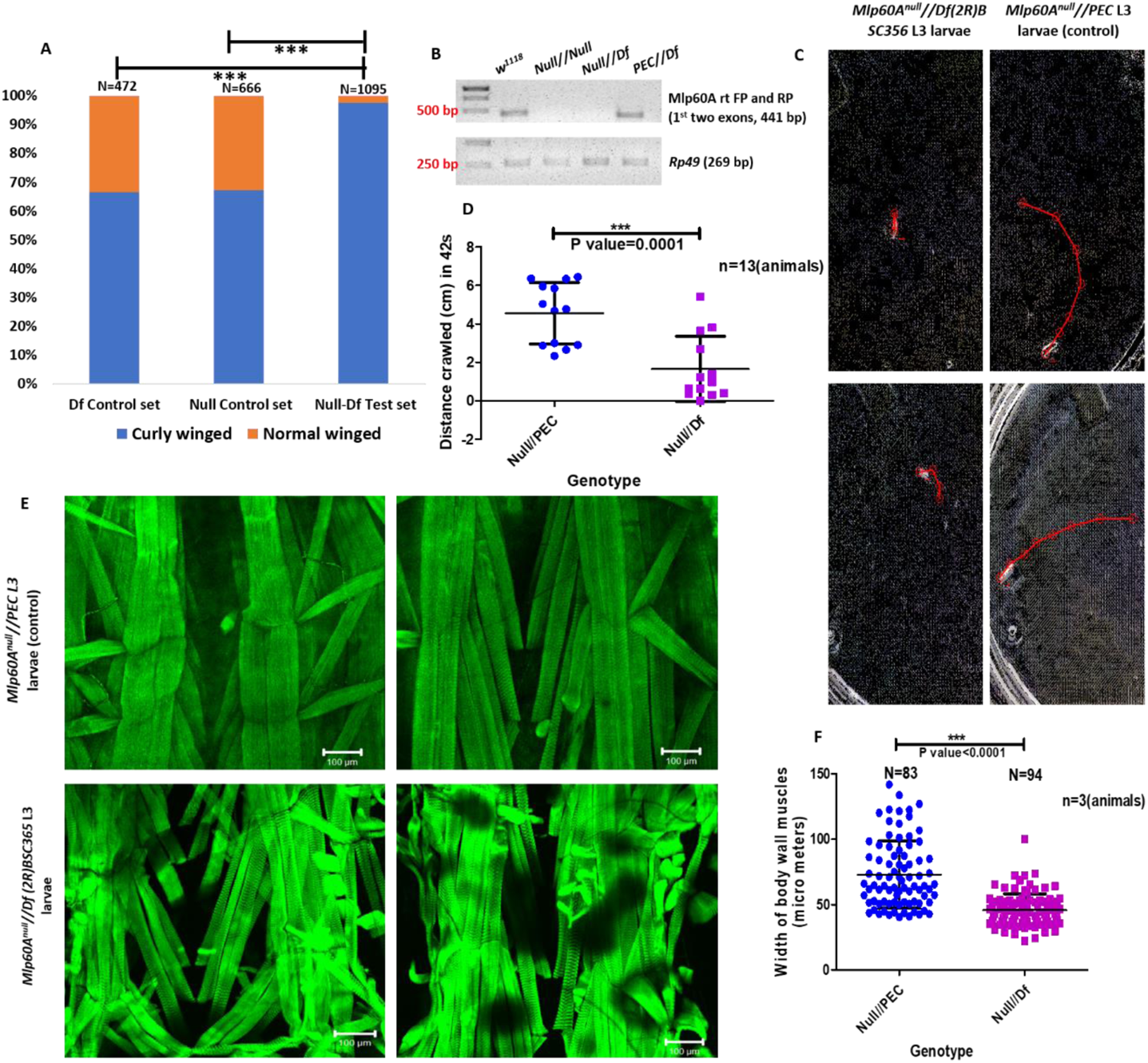
*Df(2R)BSC356-Mlp60A^null^* complementation test. (A) Shows the survival of *Df//Mlp60A^null^* trans-heterozygotes. The survival percentage of these individuals was significantly lesser than that which would be expected following normal mendelian genetics, as seen in the control. For statistical (Chi Square Test) analysis, P value<0.0001, Chi Square Value (CSV)=43.464 at degree of freedom (df)=1. Y-axis shows the percentage of flies, X-axis shows the sets of flies (B) Shows the absence of a PCR product from a reaction carried out with primers designed in the *Mlp60A* region, thus molecularly confirming the deficiency chromosome. (C) Shows representative traces of two larvae of each of the tested genotypes. (D) Quantification of the crawling ability of *Mlp60A^null^//Df(2R)BSC356* L3 larvae. (E) Body wall muscle defects seen in the *Mlp60A^null^//Df(2R)BSC356* L3 larvae. (F) Quantification of the width of body wall muscles in *Mlp60A^null^//Df(2R)BSC356* L3 larvae.

**Figure S8:**
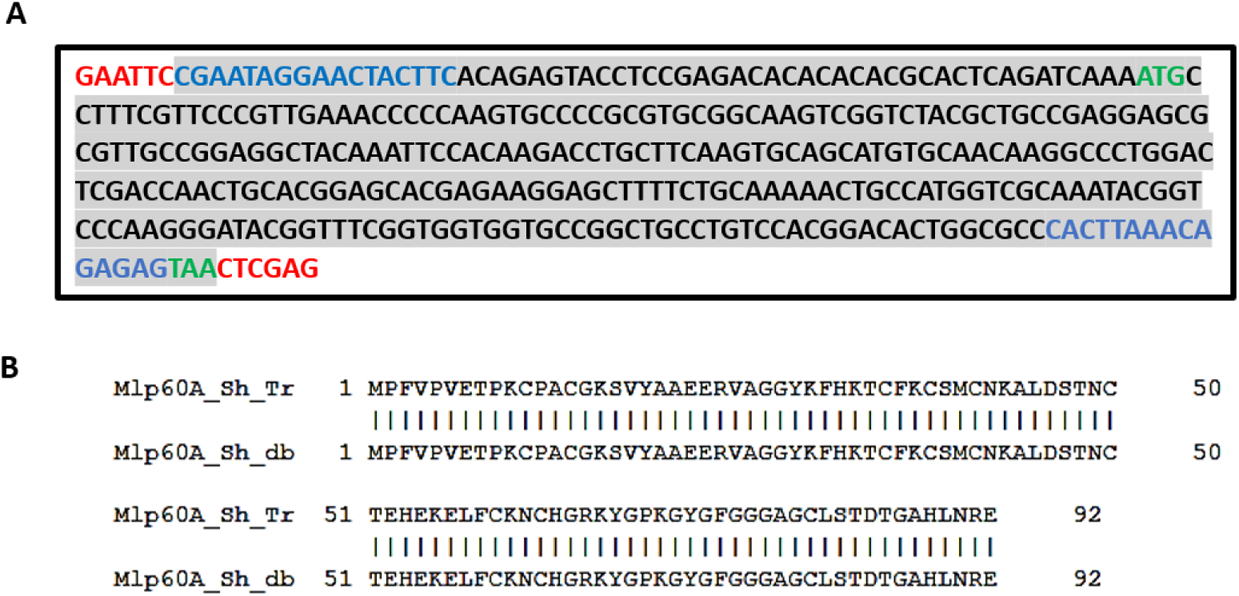
Verification of the sequence of *Mlp60A-short* CDS cloned within the *pUASt-attB* vector. (A) Shows the *Mlp60A-short* CDS with *EcoR1* and *Xho1* restriction sites. (B) Shows the alignment of the translated *Mlp60A-short* CDS with the protein sequence of this isoform, from the database.

**Figure S9:**
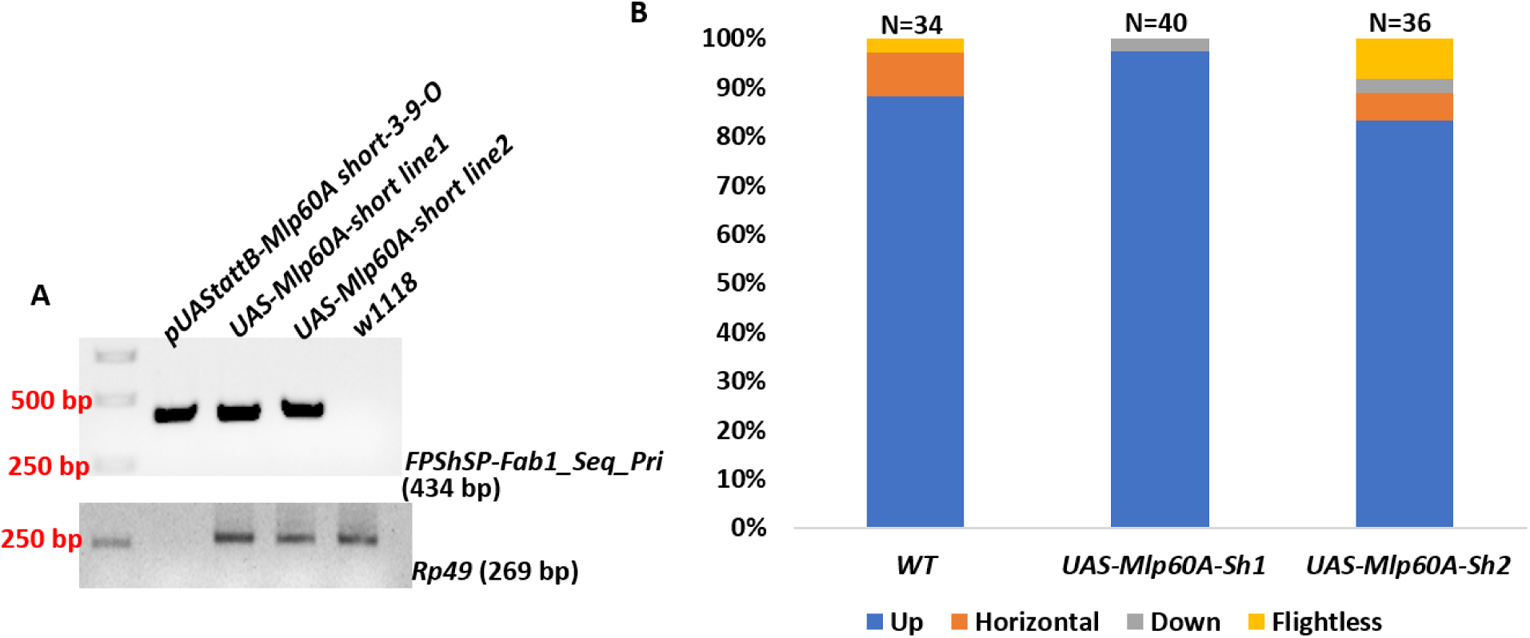
Verification of *pUASt-attB-UAS-Mlp60A-short* transgenic lines. (A) Shows the presence of the transgenic construct in the genomic DNA of the transgenic lines. (B) Shows the flight ability of the flies homozygous for the *pUASt-attB-UAS-Mlp60A-short* construct bearing chromosomes. Y-axis shows the percentage of flies, X-axis shows the genotype of flies.

**Figure S10:**
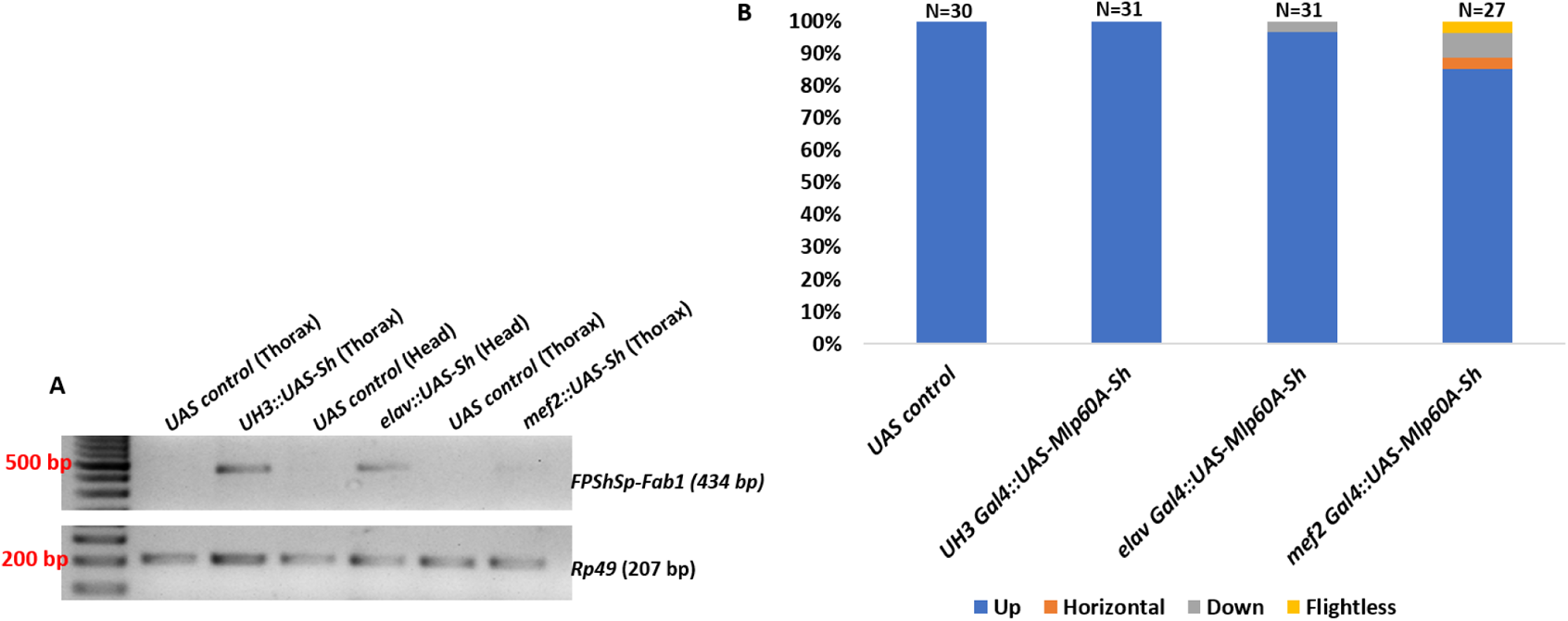
Transgenic over-expression of *Mlp60A-short* isoform, through the *UAS-Gal4* system. (A) Shows the presence/absence of the transgenic *Mlp60A-short*. (B) Shows the flight ability of the flies arising from different sets of over-expression crosses. Y-axis shows the percentage of flies, X-axis shows the genotype of flies.

**Figure S11:**
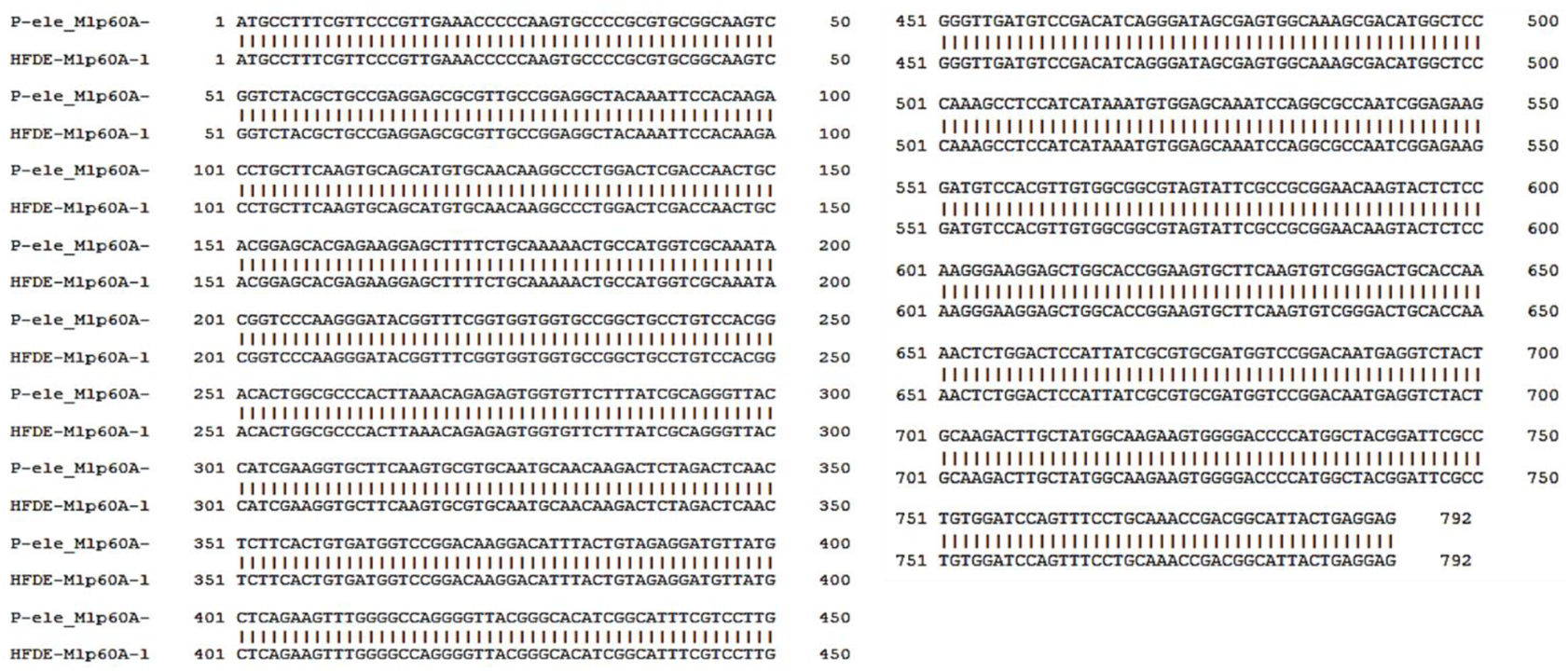
Alignment of *Mlp60A-long* mature transcripts encoded by *P-ele* and *HFDE* alleles. These are not the full-length sequences of the *Mlp60A-long* transcripts encoded by these alleles, but the sequences amplified by the primer pair *FPShSp-RPLoSp*, shown in Fig. 7B.

**Figure S12:**
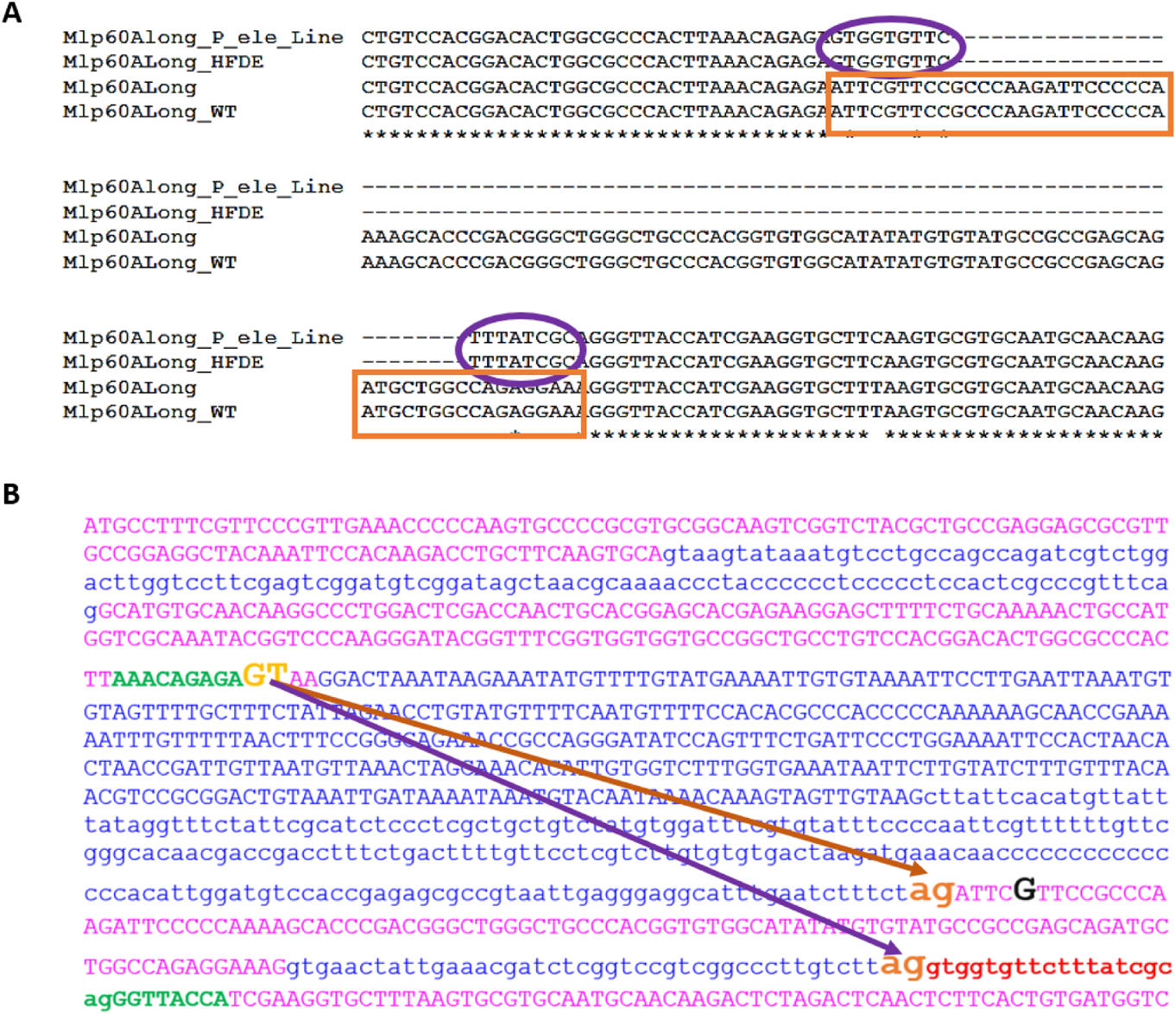
Analysis of *Mlp60A-long* transcripts encoded by *P-ele* allele and *HFDE* allele. (A) Shows a portion of the alignment between *Mlp60A^P-ele^*, *Mlp60A^HFDE^* and wild-type allele encoded *Mlp60A-long* mature transcripts, along with the database sequence as a reference. For detailed explanation see text (B) Shows the alternative splicing event occurring in the *Mlp60A^P-ele^* and *Mlp60A^HFDE^* homozygous mutants, which leads to the expression of the mutant *Mlp60A-long* mature transcript, encoded by the respective alleles. The yellow coloured ‘GT’ serves as the splice donor in the wild-type and mutant alleles. The brown line indicates the splicing event occurring in the wild type flies, leading to the expression of the wild-type *Mlp60A-long* mature transcript. The purple line indicates the alternative splicing event occurring in the *Mlp60A^P-ele^* and *Mlp60A^HFDE^* homozygous mutants which leads to the selection of an alternate splice acceptor and thus excludes the entire 3^rd^ exon sequence and inclusion of 17 bases shown in red colour (1726^th^-1742^nd^ base), in the mature transcript.

**Figure S13:**
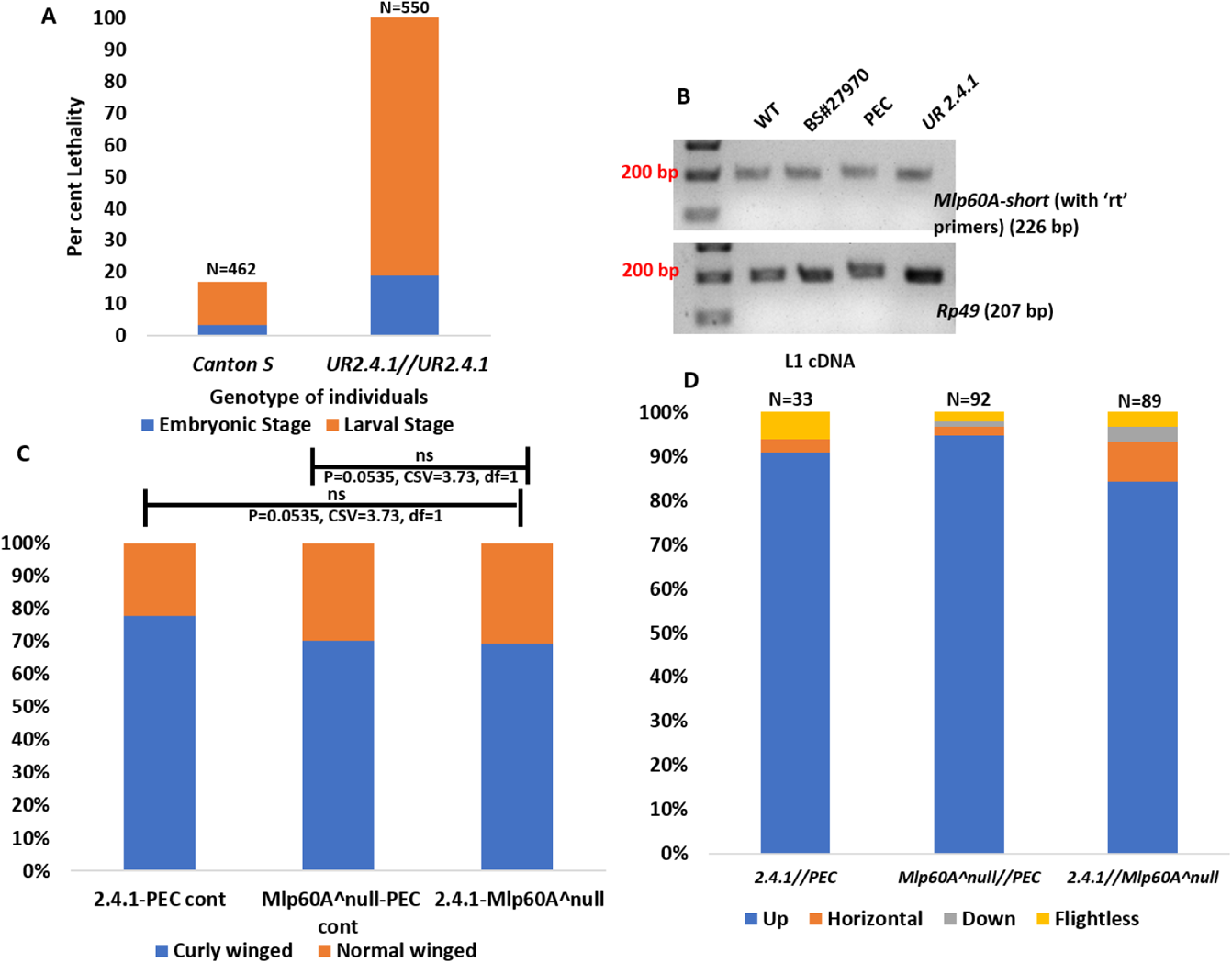
Characterization of the isolated allele *UR 2.4.1*. (A) Assessment of developmental lethality, beginning from the egg (embryo) stage. (B) Shows the *Mlp60A-short* isoform, detected through RT-PCR, from *UR 2.4.1* homozygous individuals. (C) Shows the survival percentage of *UR 2.4.1//Mlp60A^null^* trans-heterozygous flies in a complementation test between these two alleles. The survival percentage of the *Mlp60A^null^//UR 2.4.1* flies was not significantly different from that of either the *UR2.4.1//PEC* flies or that of the *Mlp60A^null^//PEC* flies (both being the control genotypes). (D) Shows flight ability of *UR 2.4.1//Mlp60A^null^* trans-heterozygous flies. Y-axis shows the percentage of flies, X-axis shows the genotype of flies.

**Table S1:**
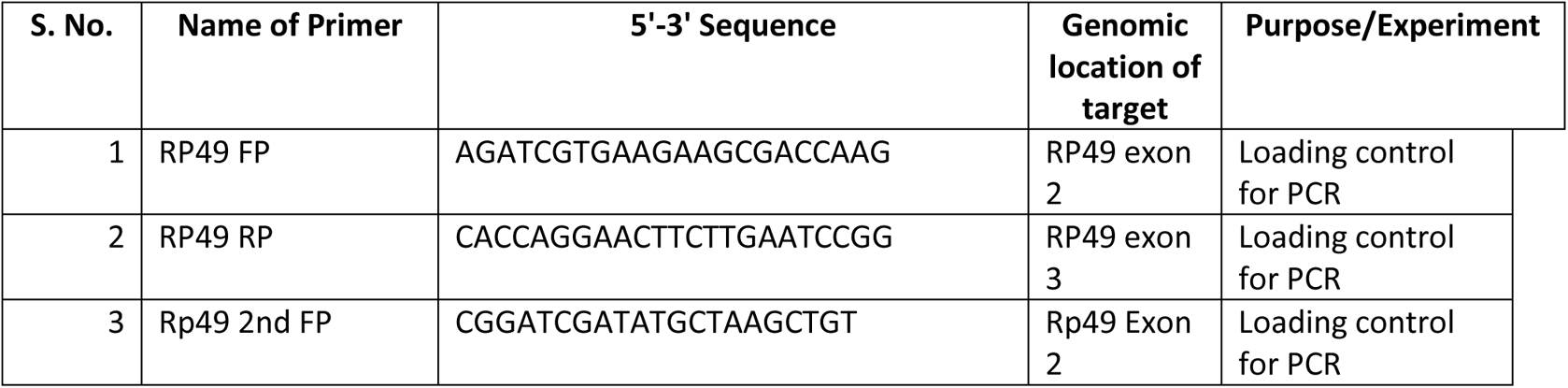
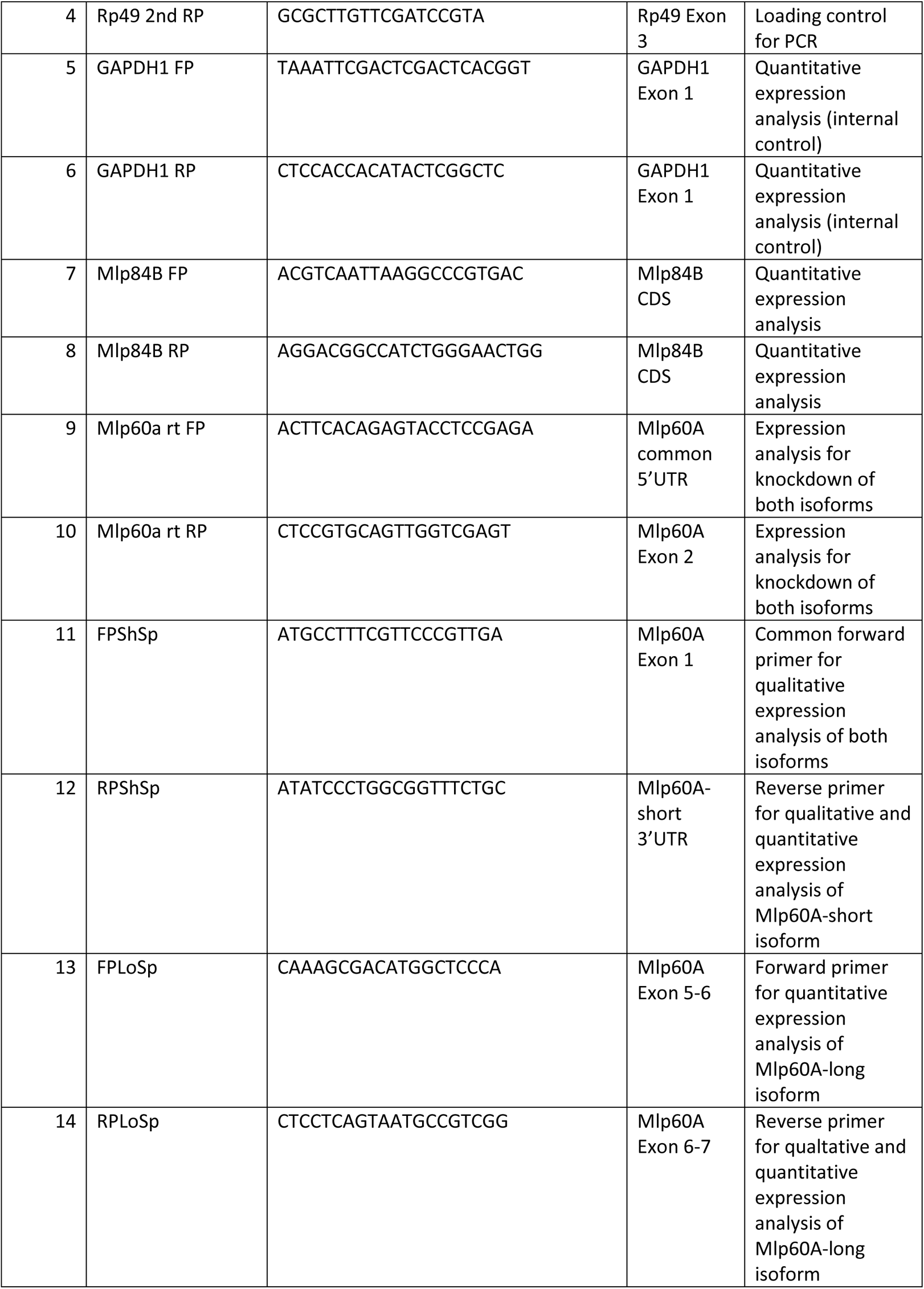
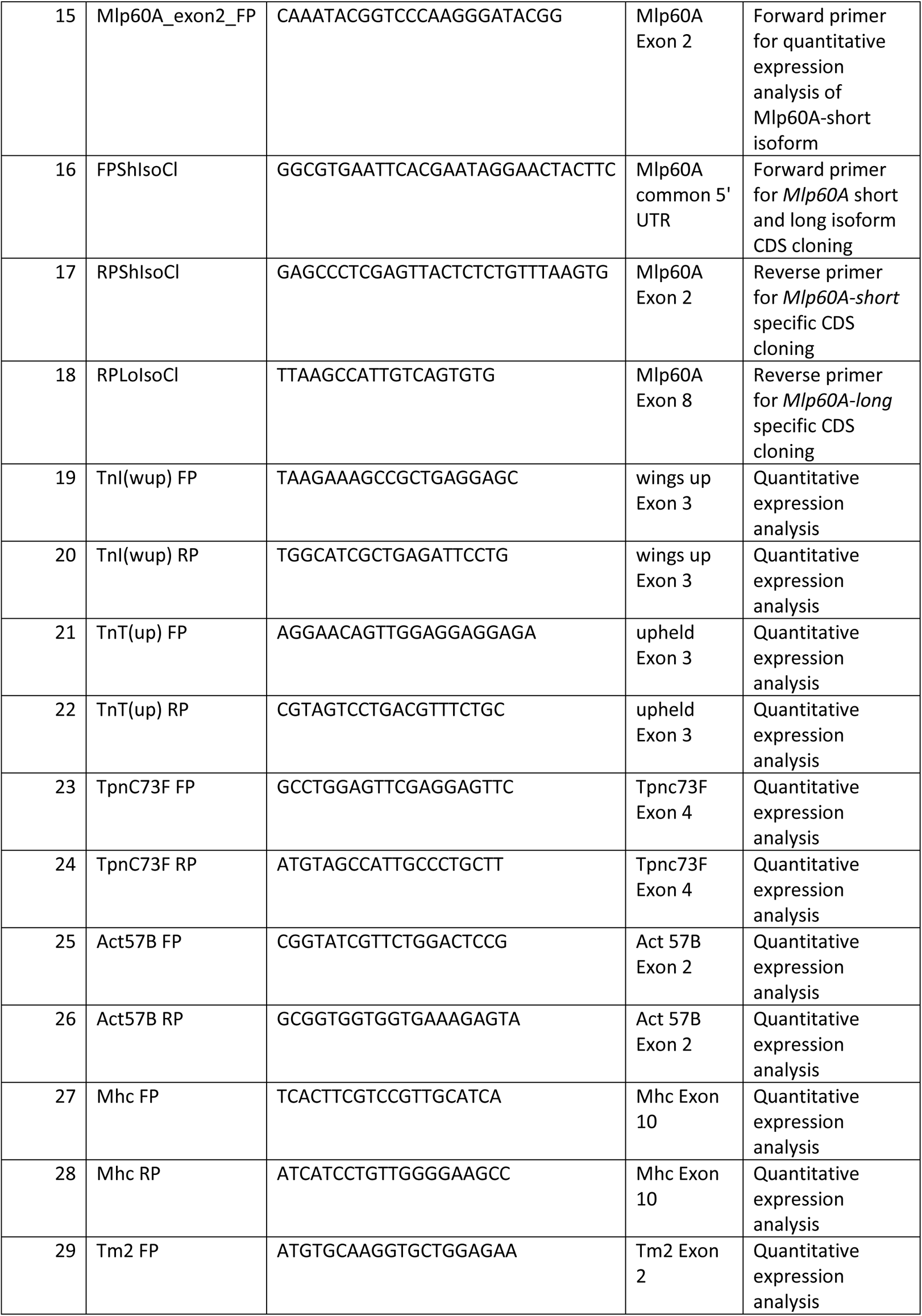
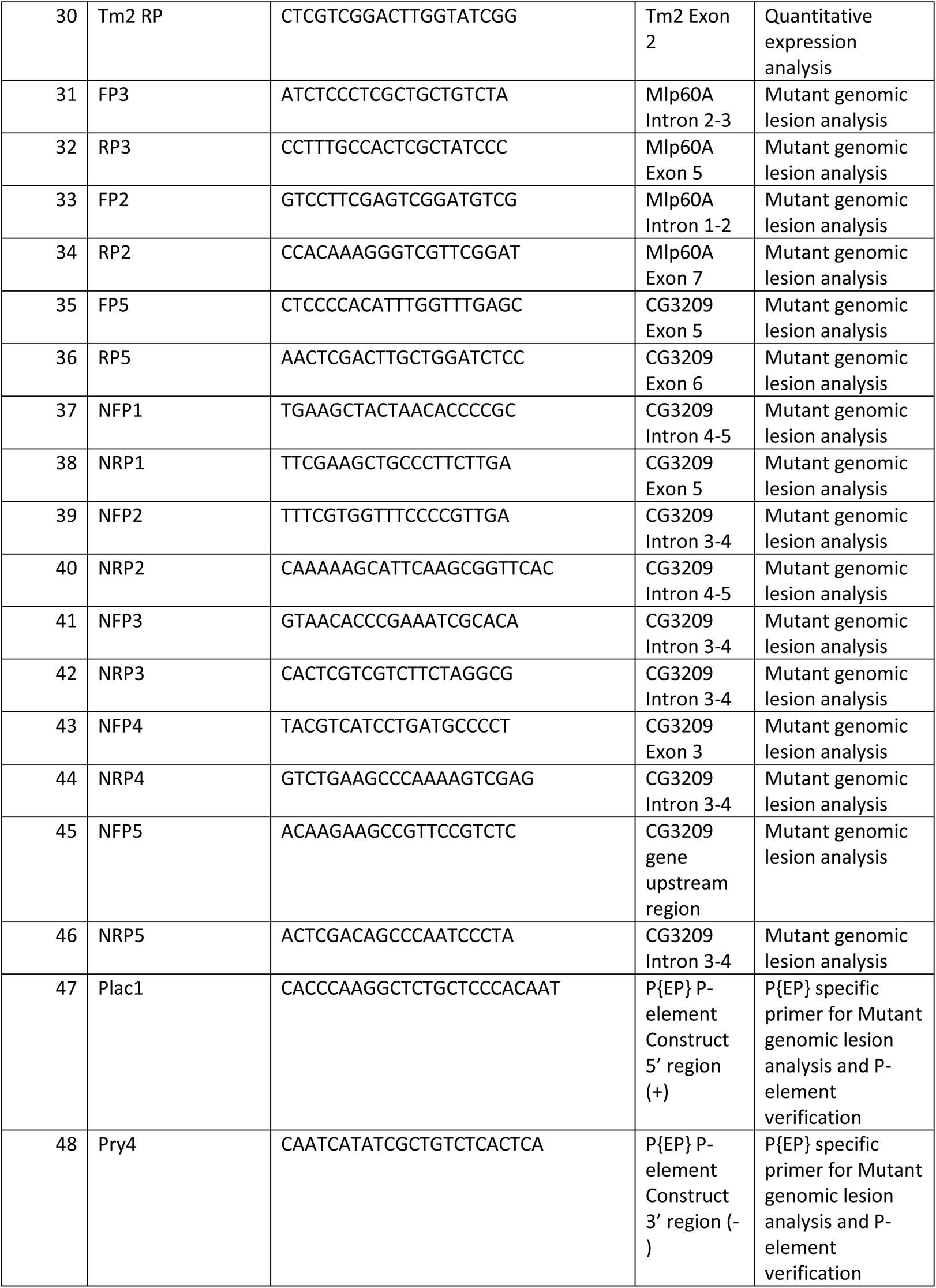
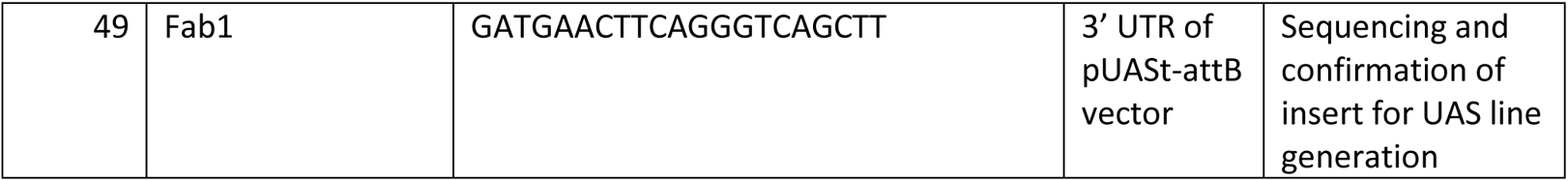
Primers used in this study.

